# Static and dynamic DNA loops form AP-1 bound activation hubs during macrophage development

**DOI:** 10.1101/142026

**Authors:** Douglas H. Phanstiel, Kevin Van Bortle, Damek Spacek, Gaelen T. Hess, Muhammad Saad Shamim, Ido Machol, Michael I. Love, Erez Lieberman Aiden, Michael C. Bassik, Michael P. Snyder

## Abstract

The three-dimensional arrangement of the human genome comprises a complex network of structural and regulatory chromatin loops important for coordinating changes in transcription during human development. To better understand the mechanisms underlying context-specific 3D chromatin structure and transcription during cellular differentiation, we generated comprehensive in situ Hi-C maps of DNA loops during human monocyte-to-macrophage differentiation. We demonstrate that dynamic looping events are regulatory rather than structural in nature and uncover widespread coordination of dynamic enhancer activity at preformed and acquired DNA loops. Enhancer-bound loop formation and enhancer-activation of preformed loops represent two distinct modes of regulation that together form multi-loop activation hubs at key macrophage genes. Activation hubs connect 3.4 enhancers per promoter and exhibit a strong enrichment for Activator Protein 1 (AP-1) binding events, suggesting multi-loop activation hubs driven by cell-type specific transcription factors may represent an important class of regulatory chromatin structures for the spatiotemporal control of transcription.

**HIGHLIGHTS:**

- High resolution and high sensitivity of loop detection via deeply sequenced in situ Hi-C experiments during monocyte to macrophage differentiation (> 10 billion total reads)
- Multi-loop interaction communities identified surrounding key macrophage genes.
- Multi-loop communities connect dynamic enhancers through both static and newly acquired DNA loops, forming hubs of activation
- Macrophage activation hubs are enriched for AP-1 bound long-range enhancer interactions, suggesting cell-type specific TFs drive changes in 3D structure and transcription through regulatory DNA loops

## INTRODUCTION

It is estimated that less than 2% of the human genome codes for functional proteins (Alexander et al., 2010). Scattered throughout the rest of the genome are regulatory regions that can exert control over genes hundreds of thousands of base pairs away through the formation of DNA loops. Loop-based transcriptional regulation plays a part in many biological contexts but is critically important for human development and cellular differentiation (Krijger and de Laat, 2016). Alteration of DNA loops has been implicated in a variety of developmental abnormalities and human diseases. Disruption of long-range interactions between *Hoxd* genes and dispersed enhancer-like sequences leads to abnormal digit morphology in the developing limb bud (Montavon et al., 2011) and deletion of loop anchor regions within the *Hoxa* and *Hoxc* clusters leads to homeotic transformations (Narendra et al., 2016). Microdeletions affecting 3D chromatin structures and transcription of human oncogenes have been identified in many forms of cancer (Hnisz et al., 2016).

Structurally, DNA loops are controlled by both global and local genomic elements which establish a hierarchy of chromatin organization (Dekker et al., 2013). Typically acting in cis at distances no larger than 1-2 Mb, DNA loops are confined by structures known as topologically associating domains (TADs) which demarcate boundaries of hetero- and euchromatin (Dixon et al., 2012; Nora et al., 2012). TADs themselves are partitioned into two or more nuclear compartments of similar transcriptional activity (Imakaev et al., 2012; Lieberman-Aiden et al., 2009). At a local level, loops are controlled by proteins including CCCTC-binding factor (CTCF) and members of the cohesin complex, both of which are highly enriched at loop anchors, as well as cohesin loaders and unloaders (Busslinger et al., 2017; Haarhuis et al., 2017; Heidari et al., 2014; Rao et al., 2014; Sofueva et al., 2013). While our knowledge regarding the general characteristics and mechanisms of loops is improving, much less is known regarding the scope, mechanisms, and functional significance of dynamic looping events during biological processes such as cellular differentiation.

Rapidly evolving DNA sequencing-based technologies have provided increasingly comprehensive views of DNA looping in human cells and are improving our ability to address these salient questions. At their core, all of these technologies apply the same basic paradigm: DNA-DNA interactions are fixed using chemical cross-linking, fragmented by physical or enzymatic methods, and ligated together forming chimeric DNA sequences harboring DNA from each of the two interacting regions. Frequencies of ligation serve as a proxy for in vivo interaction frequencies allowing for the inference of DNA loops, TADs, nuclear compartments, and other chromatin structures. There are a wide variety, and ever increasing, number of different variations on this approach; however, the current methodologies can be classified into two categories; genome-wide and targeted (Davies et al., 2017). Hi-C is a genome-wide approach to detect contact frequencies between all mappable regions of the human genome. Application of this approach has revealed extensive chromatin reorganization, both within stable chromatin domains and in higher order compartment localization, during ES cell differentiation and across human tissues and cell-types (Dixon et al., 2015; Schmitt et al., 2016). However, the sequencing depth required to achieve high resolution maps of contact frequency as well as the high number of nonspecific ligation events have made the identification of loops using Hi-C problematic.

To circumvent these issues a number of approaches have been developed that target small fractions of the genome, thus reducing the sequencing burden required to achieve high resolution (Dostie et al., 2006; Fullwood et al., 2009; Mumbach et al., 2016). Application of one such approach, targeted chromosome conformation capture approach (5C), which interrogates small contiguous genomic stretches, identified examples of short-range non-CTCF loops that change in differentiating mouse embryonic stem cells at specific regulatory genes (Phillips-Cremins et al., 2013), and which to varying degrees are dynamically restored during somatic cell reprogramming (Beagan et al., 2016). Chromatin Interaction Analysis by Paired-End Tag Sequencing (ChIA-PET), an approach capable of investigating interactions between genomic regions bound by specific proteins, further provides examples of dynamic enhancer and promoter interactions between ES cells and differentiated B lymphocytes (Kieffer-Kwon et al., 2013). Although these studies suggest that differentiation-associated activation and repression of specific genes is accomplished through changes in chromatin architecture, such targeted approaches are unsuitable for identifying genome-wide intra-chromosomal interactions that are likely important for cellular differentiation. Moreover, by interrogating only small interspersed regions of the genome, approaches such as ChIA-PET, capture Hi-C, and HiChIP may not be able to distinguish differences in local chromatin compaction from true DNA loops (Rao et al., 2014).

The introduction of a new genome-wide approach, in situ Hi-C, in combination with the continually decreasing cost of genomic sequencing, has allowed for the unbiased genome-wide detection of DNA loops in human cells. Nuclear proximity ligation increases the efficiency of in situ HiC over previous “dilution” Hi-C protocols by reducing random ligation events, enabling higher resolution and detailed mapping of DNA loops at currently achievable sequencing depths. Importantly, the comprehensive nature of in situ Hi-C allows comparison of interaction frequencies to local backgrounds, something not possible with targeted methods like ChIA-PET, capture Hi-C, and HiChIP, producing quantifiable improvements in accuracy of loop detection (Rao et al., 2014). While multi-cell comparison of high resolution in situ Hi-C maps has identified thousands of DNA loops that are preserved across diverse cell types (Rao et al., 2014), examples of cell-type specific looping events also support a role for dynamic genome architecture regulating specific genes. However, this method has not been broadly applied to study dynamic looping in the context of human development and cellular differentiation.

Macrophages are phagocytic cells of the innate immune system that represent one of the body’s first defenses against invading pathogens. Macrophage-mediated inflammation has been identified as a key driver of multiple human disorders and diseases including atherosclerosis, diabetes, and cancer (Chawla et al., 2011; Moore et al., 2013; Noy and Pollard, 2014; Ostuni et al., 2015). The differentiation of monocytic precursors into mature macrophages is well characterized and key transcriptional regulators of this process have been identified including SPI1, MAFB, and AP-1 (Kelly et al., 2000; Rosa et al., 2007; Valledor et al., 1998). However, the role of DNA looping in this process remains completely unexplored. Treatment of the monocytic leukemia cell line, THP-1, with phorbol myristate acetate (PMA), is a widely used model for studying monocyte-macrophage differentiation and provides an ideal system for studying the regulatory dynamics controlled via long-range interactions (Daigneault et al., 2010). First, THP-1 cells grow as a largely homogenous cell population with a near-diploid genetic background lacking major cytogenetic rearrangements typical of most established cell lines (Odero et al., 2000). Second, they exhibit high functional similarity to in vivo monocytes including the ability to differentiate into extremely pure populations of macrophages (Kouno et al., 2013). And finally, because these cells renew indefinitely, unlimited experiments can be performed on these same cells eliminating variability introduced by genetic differences.

Here, we apply in situ Hi-C in combination with other genomic methodologies to profile global changes in DNA looping events during the differentiation of human monocytes into macrophages. We show that the transcriptional dynamics of differentiating THP-1 cells are accompanied by significant changes in long-range DNA loops at key regulatory genes known to be important for macrophage development and function. Intersection with histone modification profiles reveals that transcriptional regulation is accompanied by both gained and pre-formed chromatin loops that acquire enhancer activity during differentiation. These gained and enhancer-activated loops form multiple-loop regulatory communities that connect single promoters to multiple distal enhancers. Strong enrichment of AP-1 binding sites is observed at the promoter-distal ends of these gained and activated loops. Taken together these results suggest that the formation of multiloop activation hubs increases the local concentration of AP-1 and transcriptional activation machinery at the promoters of key regulatory genes promoting increased transcription during macrophage development. AP-1 activation hubs are similar in nature to specific, previously characterized multi-loop structures, such as the developmentally regulated beta globin locus, suggesting that activation hubs may represent a common mechanism for controlling the spatiotemporal expression of distinct regulatory genes important for other developmental contexts.

## RESULTS

### High resolution maps of DNA structure in human monocytes and macrophages

To determine how chromatin structure changes during cellular differentiation, we applied in situ Hi-C to THP-1 cells before and after exposure to PMA, generating high resolution genome-wide maps of DNA-DNA interaction frequencies in both THP-1 monocytes and THP-1 derived macrophages. Sequencing to a depth of greater than 5 billion reads per sample, we generated high quality normalized contact maps with a bin resolution of 10 kb (Fig. 1A). To better understand the nature of DNA looping during PMA-induced differentiation we generated quantitative data sets that mapped changes in transcript abundance (RNA-seq), chromatin accessibility (ATAC-seq), CTCF occupancy, and H3K27 acetylation levels (ChIP-seq), a histone modification associated with active enhancers (Fig. 1B; (Creyghton et al., 2010)). DNA loops were determined by identifying contact frequencies (pixels) that are significantly enriched compared to local background levels using a method developed by Rao et al (see supplemental methods; (Rao et al., 2014)). We identified 16067 and 16335 long-range interactions in untreated and treated THP-1 cells respectively (**Tables S1,S2**). Hi-C data sets were also processed by Juicer yielding very similar results (Fig. S1; (Durand et al., 2016)). Quality of loop calls can be assessed by aggregating Hi-C signal in the pixels surrounding all detected loops using a method called Aggregate Peak Analysis (APA) (Rao et al., 2014)). High quality DNA loops calls are characterized by APA plots with intense center pixels surrounded by less intense pixels and these plots can be scored by dividing the value of the center pixel by the median of pixels in the bottom right quadrant. Application of the APA method to our loop calls for both untreated and treated cells revealed extremely high scoring APA plots (2.66 and 3.2) highlighting the accuracy of the loops detected (Fig. 1C-D).

**Figure 1:**
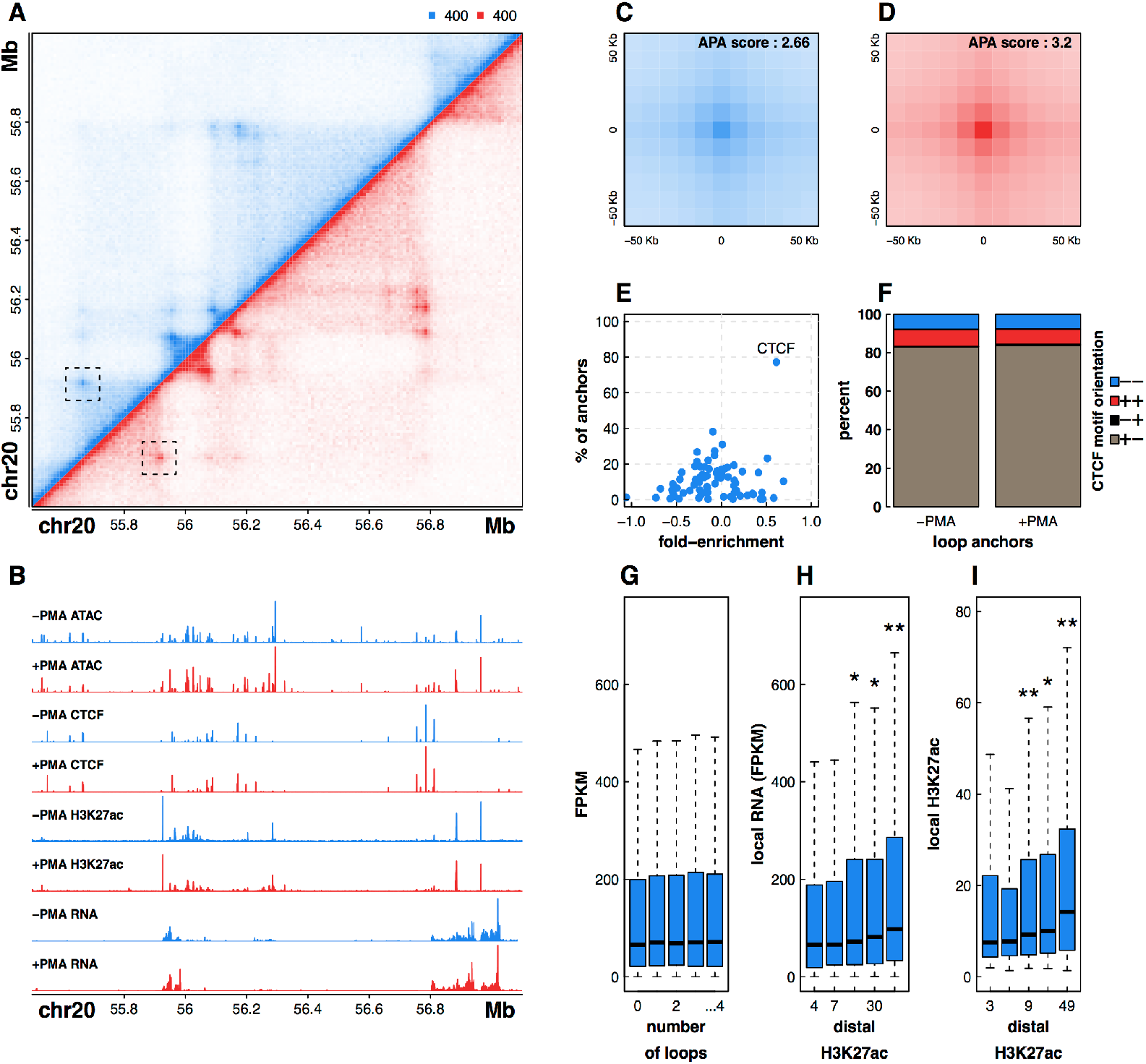
Integrative genomic profiling reveals principles of DNA looping in untreated and +PMA treated THP-1 cells. (**A**) Hi-C contact matrix depicting normalized contact frequencies between loci on chromosome 20 for untreated THP-1 cells (blue, top left) and PMA treated THP-1 cells (red, bottom right). One of the loops detected in both untreated and treated cells is highlighted. (**B**) ATAC-seq, ChIP-seq, and RNA seq signal tracks corresponding to the Hi-C region depicted above. APA plots showing aggregated signal across all loops in both untreated (**C**) and PMA treated THP-1 cells (**D**). (**E**) Enrichment of CTCF motifs in loop anchors of untreated THP-1 cells. (**F**) Stacked bar plot depicting CTCF motif orientations at loop anchors as a percentage of all loops that contain a single CTCF bound peak at each anchor. (**G**) RNA FPKM values of genes binned by the number of loops that connect to their promoter. (**H**) RNA FPKM values of genes binned by the histone H3K27 acetylation signal at the promoter-distal end of a loop. (**I**) H3K27 acetylation signal binned by the histone H3K27 acetylation signal at the other end of a loop reveals correlated histone H3K27 acetylation at loop anchors. Significant differences (p < 0.05, Wilcox Rank Sum Test) vs subset to the immediate left or all subsets to the left are indicated by one of two asterisks respectively. See also Figures S1 and S2.

To further assess the quality of our data we examined transcription factor binding motifs in the anchor regions of our detected loops. DNA motif enrichment analysis identified the CTCF target sequence at nearly 80% of all loop anchors (Fig. 1E). Long-range interactions between CTCF target sequences were recently shown to possess strong orientation bias, with most DNA loops connecting two inward facing CTCF elements pointing in the forward and reverse orientations (Rao et al., 2014). Inversion of either CTCF motif is sufficient to disrupt the directionality of DNA-looping interactions, suggesting CTCF plays a significant role in establishing chromatin architecture (Guo et al., 2015). Consistent with previous reports, our analysis of CTCF motif orientation at THP-1 loop anchors revealed a strong inward facing CTCF orientation bias at loop ends before and after differentiation (Fig. 1F). The predominance of CTCF, which is thought to mediate long-range interactions via recruitment of the cohesin complex (Merkenschlager and Nora, 2016), and orientation bias in CTCF binding at loop anchors, further serve to validate the accuracy and quality of our chromatin interactome map in THP-1 cells. Despite this strong enrichment for CTCF motifs, we also identify numerous clear examples of CTCF-independent looping (Fig. S2), raising critical questions regarding the mechanisms driving the formation and maintenance of DNA loops in human cells and highlighting the need for truly genome-wide interaction mapping approaches such as in situ Hi-C.

To explore the regulatory nature of DNA looping in monocytes and macrophages we next intersected our loop calls with histone H3K27 acetylation signal and transcript abundance. Surprisingly, comparison of relative transcript level and the number of long-range interactions that a promoter is connected to reveals no significant relationship between gene activity and the number of DNA loops (Fig. 1G). However, we find that both transcription and local H3K27 acetylation levels are significantly higher for regions that are connected to distal DNA elements with elevated H3K27 acetylation, implying active enhancer-gene interactions (Fig. 1H-I). Overall, this result suggests that the activity and chromatin context at distal regulatory element connections, rather than the mere presence or number of long-range interactions, is a major determinant of gene expression and perhaps an important element controlling dynamic transcription of regulatory genes during cellular differentiation.

### Distal regulation of macrophage-specific transcription is associated with both dynamic and pre-formed DNA loops

We next leveraged our longitudinal in situ Hi-C experiments to determine whether differentiation of THP-1 cells is accompanied by dynamic looping events. Using a method that explicily tests for changes relative to local background, we identified 217 significant changes in DNA looping events in THP-1 cells, with a loss in 33 monocyte-enriched interactions and a gain in 184 macrophage-enriched interactions after treatment with PMA (DESeq2, p < 0.001; **Supplemental Methods, Table S3**). APA analysis of differential looping events illustrates both the reproducibility of dynamic interactions across biological replicates and the significant difference in contact frequencies before and after differentiation of THP-1 cells, suggesting our stringent differential loop calling identifies loops that are entirely lost or gained during macrophage development (Fig. 2A-C). Examples of loops that were static, lost, or gained during differentiation are shown in Figure 2D-F. Overall, we find that loops gained during differentiation are more abundant (184 vs 33) and overlap more genes (88 vs 8) than loops lost during differentiation. Inspection of both APA plots (Fig. 2B-C) and individual loops further support this bias in dynamic looping, altogether suggesting that loop formation as opposed to loop disruption may play a broader role in macrophage development.

**Figure 2:**
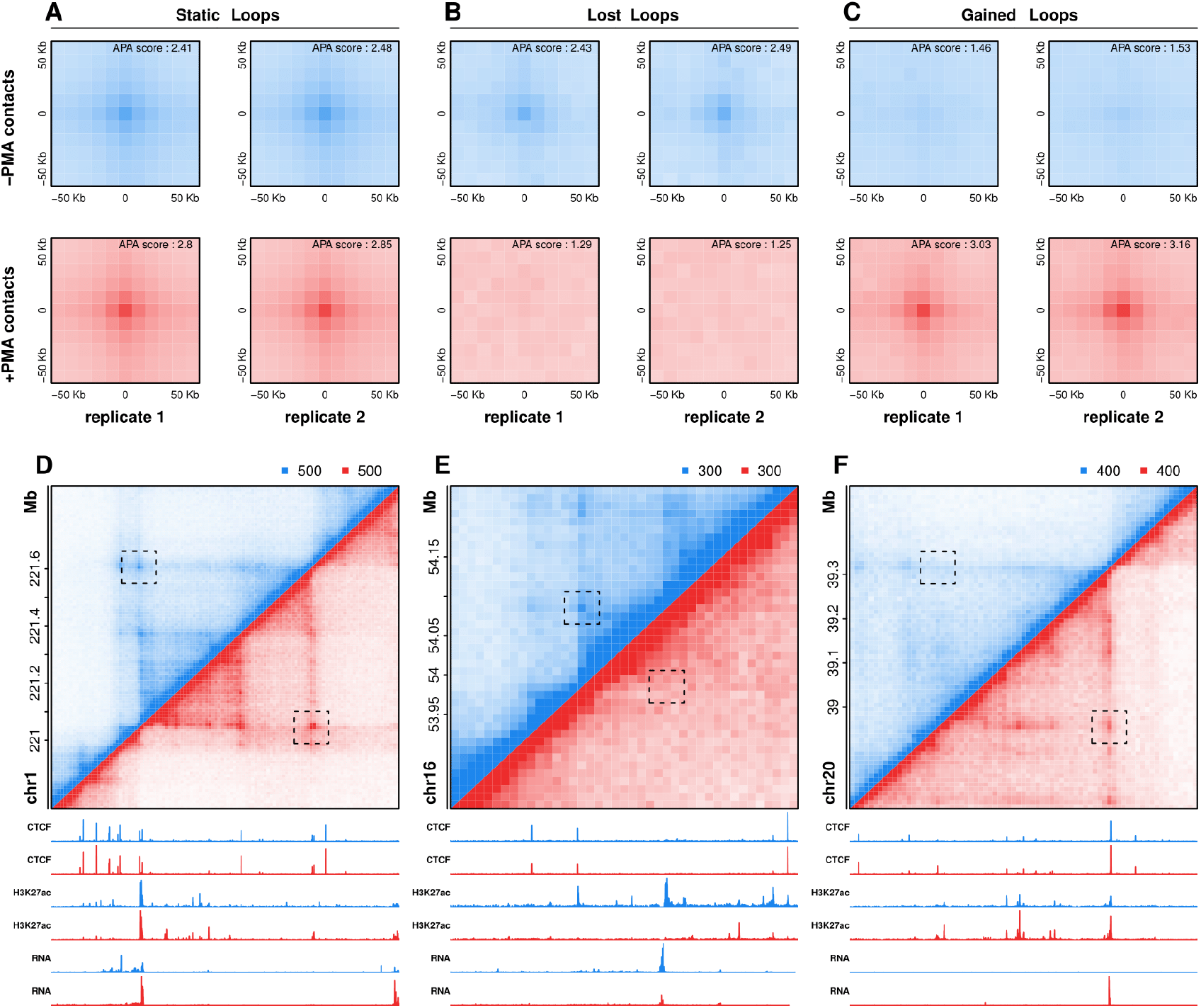
Detection and visualization of differential looping events during differentiation. APA plots for loops that are static (**A**), lost (**B**), or are gained during PMA-induced differentiation of THP-1 cells. Individual plots for each biological replicate and condition are shown. Hi-C contract matrices, ChIP-seq signal tracks, RNA-seq signal tracks, and genes are shown for regions surrounding an examples of static (**D**), lost (**E**), or gained (**F**) loops.

Visual inspection of the data indicated that both static and dynamic loops link gene promoters to distal regulatory elements marked by changes in H3K27 acetylation, suggesting that in addition to dynamic looping events, preformed loops may also play an important role in distal regulation of transcription during differentiation. To investigate this further, we categorically divided our loops into five subsets; (1) loops that disappeared during differentiation (‘lost’), (2) loops that did not change significantly during differentiation but contained an anchor region that harbored a decreasing promoter-distal H3K27ac peak (‘deactivated’), (3) loops that did not change and that did not overlap any dynamic promoter-distal H3K27ac peaks (static’), (4) loops that did not change during differentiation but contained an anchor region that harbored an increasing promoter-distal H3K27ac peak (‘activated’), and (5) loops that are acquired during differentiation (‘gained’) (Fig. 3A). We next considered how genes whose promoters overlapped anchors of each of these loop sets changed during differentiation (Fig 3B-C). For deactivated and activated enhancer-loop sets, only genes at the distal end of the dynamic enhancer were considered. Only eight genes were found at lost loops and they showed no significant difference in expression when compared to static loops. However, deactivated loops connected regions with decreased H3K27 acetylation to 72 genes whose expression significantly decreased during differentiation. In contrast, activated and gained loops overlapped 455 and 88 genes, respectively, that increased in expression during differentiation. Overall, these results support two modes of distal gene regulation during THP-1 differentiation. In the first mode, new loop formation connects distal regulatory elements to the promoters of key regulatory genes that are dynamically expressed during differentiation. In the second mode, changes in enhancer activity control the expression of target genes through stable chromatin loops that do not change during differentiation.

**Figure 3:**
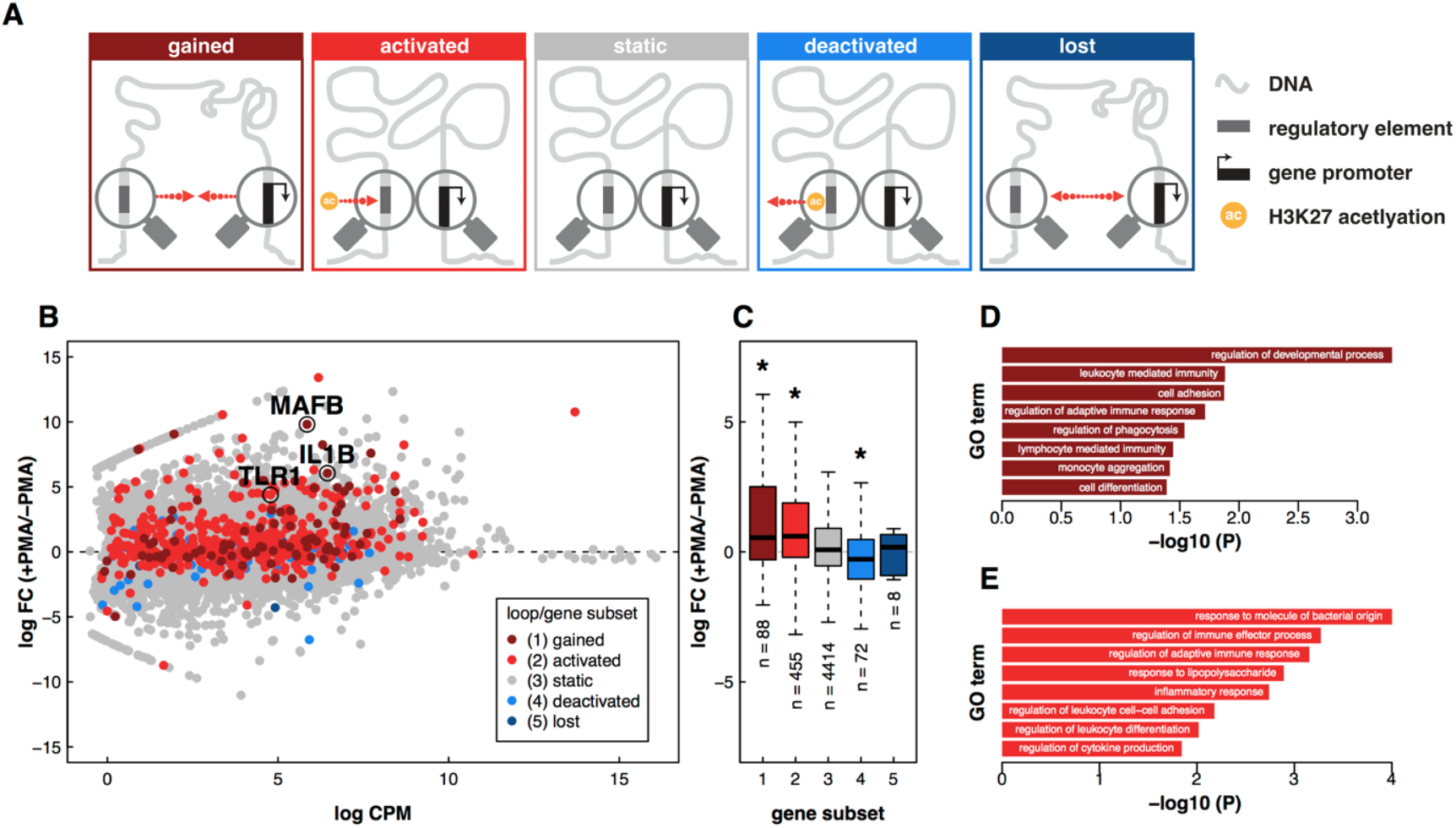
Expression and function of genes correlate with dynamic loop type and distal chromatin state. (**A**) Schematic depictions of five distinct loop classes. Red arrows indicate direction of change during PMA-induced differentiation of THP-1 cells. ‘Ac’ refers to a change in H3K27 acetylation as detected by ChlP-seq. (**B**) Scatter plot portraying mean counts per million vs log fold change for all transcripts measured by RNA-seq. Genes are colored according to the class of loop found at their promoter. (**C**) Boxplot depicting RNA FPKM values as a function of gene subset. Selected Gene Ontologies that are enriched in genes at gained (**D**) and activated (**E**) loop anchors.

Genes found at the anchors of gained and activated loops included key regulatory genes required for macrophage function and development. The genes include; (1) Interleukin 1 beta (*IL1β*), a canonical proinflammatory cytokine whose activation via proteolytic cleavage initiates widespread signaling and transcriptional changes in macrophages and neighboring cells, (2) V-maf musculoaponeurotic fibrosarcoma oncogene homolog B (*MAFB*), an AP-1 family transcription factor and essential regulator of differentiation and selfrenewal during haematopoiesis, and (3) Toll-Like Receptor 1 (*TLR1*), a cell surface membrane receptor critical for recognition of foreign pathogens and stimulation of innate immune response. Gene ontology (GO) enrichment analysis revealed that genes overlapping gained loop anchors are enriched for biological processes related to macrophage function, such as cell adhesion, phagocytosis, and immune response (Fisher’s Exact Test, p < .05; Fig. 3D, **Table S4**). GO analysis of genes associated with activated loops revealed enrichment of macrophage-related functions including response to molecule of bacterial origin, regulation of leukocyte differentiation, and inflammatory response (Fig. 3D, **Table S5**). The correlation between gene expression, enhancer activity and dynamic looping events at these macrophage-related genes together suggests that cell type specific transcription is controlled through both re-wiring of enhancer-promoter interactions and through increasing or decreasing enhancer activities at the promoter-distal end of pre-formed loops.

### Loops gained during differentiation are regulatory rather than structural in nature

One of the primary proposed functions of DNA loops is to bring distal regulatory elements such as enhancers into close physical proximity of their target genes to drive increased expression. However, many loops do not overlap gene promoters and are thought to play a structural, as opposed to gene regulatory, role (Ong and Corces, 2014). One proposed role for non-enhancer-promoter loops is the creation of insulated neighborhoods which facilitate stochastic interactions between enhancers and promoters within them (Dowen et al., 2014). However, it is currently unclear whether loops formed during differentiation are mediating direct enhancer-promoter interactions or whether they are more structural in nature. To explore this question, we used promoter-distal H3K27 acetylation as a proxy for enhancer activity to examine the degree to which static and dynamic loops regulated enhancer-promoter interactions during PMA-induced differentiation.

Loops gained during differentiation were enriched for both H3K27 acetylation and gene promoters compared to static loops (Fisher’s Exact Test, p < 0.01; Fig 4A-B). Amazingly, 90% of loops gained during differentiation connected two ‘regulatory elements’ (i.e. an enhancer to another enhancer, a promoter to another promoter, or an enhancer to a promoter) compared to only 42% of static loops. There was an especially strong enrichment for loops with H3K27 acetylation peaks at both ends (Fisher’s Exact Test, p = 1.4 × 10^−27^; Fig 4A). Greater than 60% of gained loops contained H3K27 acetylation sites at both anchors, a 2.4 fold increase compared to static loops (Fig 4A). In agreement with these results, fold changes of H3K27ac signal at loop anchors were positively correlated with changes in DNA looping (Fig 4C). Moreover, enhancers at dynamic loop anchors showed increased cell-type specificity compared to those at static loop anchors, in agreement with previous findings that differential looping plays a role in establishing cell-type specific expression patterns (Fig 4D; (Rao et al., 2014; Roadmap Epigenomics et al., 2015; Sanyal et al., 2012; Smith et al., 2016)). These findings suggest that changes in chromatin structure during PMA-induced differentiation of THP-1 cells primarily facilitate direct connections between enhancers and promoters to regulate cell-type specific gene transcription as opposed to forming structural elements such as insulated neighborhoods.

**Figure 4:**
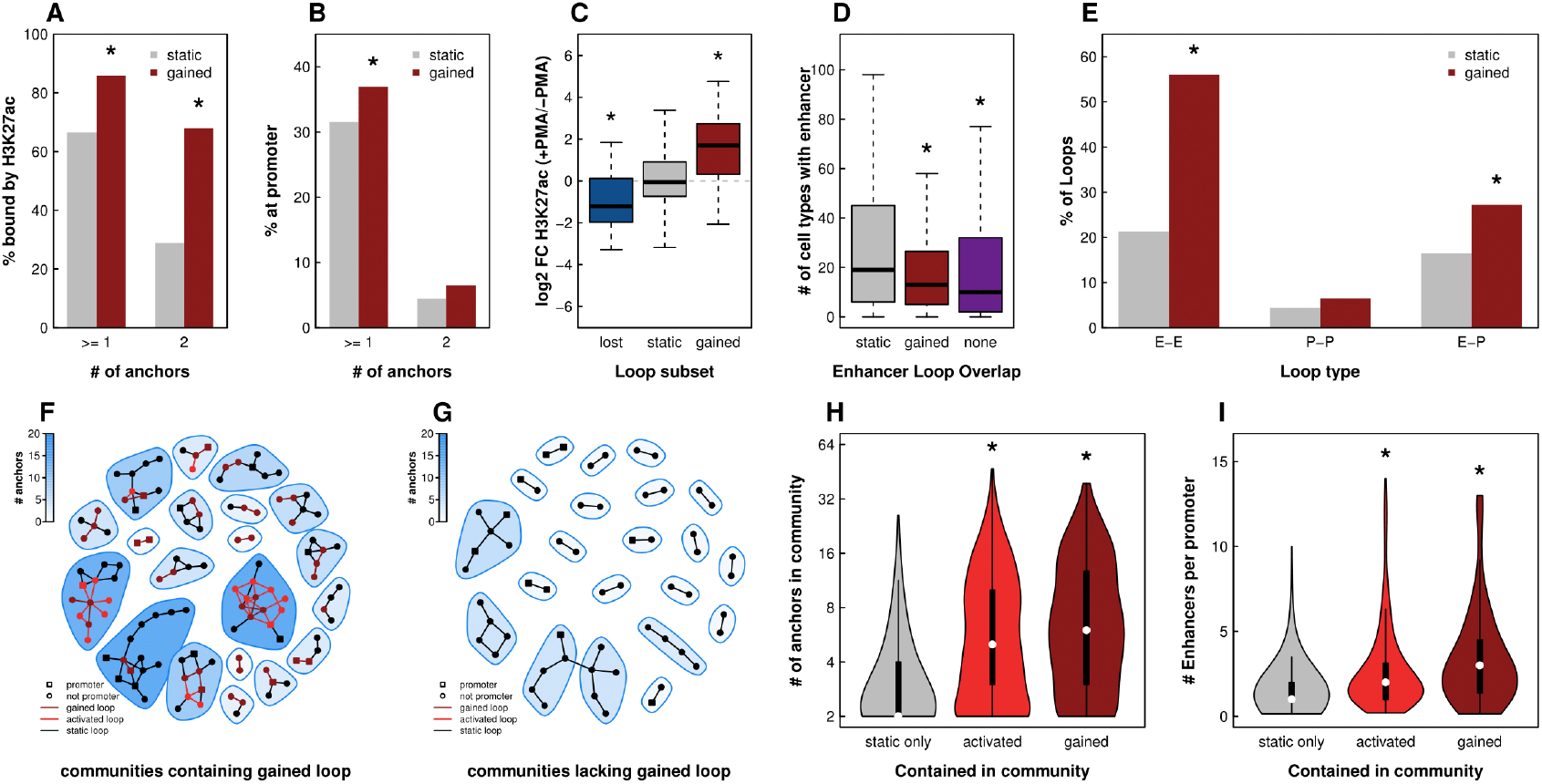
Gained and activated loops form multi-loop multi-enhancer activation hubs. (**A**) Bar plot depicting the percent of static (grey) or gained (red) loops with H3K27 acetylation peaks at the anchors. Asterisks indicate p < 10^−16^ based on Fisher’s Exact Test. (**B**) Bar plot depicting the percent of static (grey) or gained (red) loops with gene promoters at the anchors. Asterisks indicate p < 10^−4^ based on Fisher’s Exact Test. (**C**) Box plot showing the fold changes of H3K27 acetylation peaks at lost, static, and gained loop anchors. Asterisks indicate p < 10^−3^ based on Wilcox Rank Sum Test. (D) Enhancers that overlapped with static loop anchors, gained loop anchors, or no loop anchors were intersected with enhancers from 98 cell types assayed by the Roadmap Epigenomics Mapping Consortium. The number of cell types containing each enhancer is depicted as a box plot. Asterisk indicates p < 10^−4^ based on Wilcox Rank Sum Test. (**E**) Bar plot depicting percent of static (grey) or gained (red) loops that connect an enhancer to an enhancer, a promoter to a promoter, or an enhancer to a promoter. Asterisks indicate p < 10^−3^ based on Fisher’s Exact Test. 20 randomly chosen interaction communities either containing (**F**) or lacking (**G**) a gained loop are shown. Circles indicate loop anchors and lines indicate DNA loops. Colors of lines and symbols indicate type of loop. Shape of symbols indicate whether or not the loop anchor contains a gene promoter. (**H**) Violin plots showing the distribution of number of anchors per community for communities lacking gained/activated loops (static), communities containing activated loops, and communities containing gained loops. (**I**) Violin plots the distributions of enhancers to promoter ratio for communities lacking gained/activated loops (static), communities containing activated loops, and communities containing gained loops. See also Figure S3.

### Formation of mutli-loop activation hubs during macrophage development

We next explored the combinations of regulatory elements that are brought together by loops formed during differentiation (Fig 4E). Surprisingly, despite an enrichment for enhancer-promoter interactions (1.7 fold compared to static loops), direct enhancer-promoter interactions accounted for only 27% of gained loops (Fig 4E). The majority (56%) of gained loops connected an enhancer to another enhancer, a 2.6-fold enrichment compared to static loops. While the functional significance of enhancer-promoter loops is well established, the role of enhancer-enhancer interactions is less obvious.

One possible explanation for the large proportion of enhancer-enhancer loops is the presence of interaction hubs involving a single promoter and multiple enhancers that all interact with each other. A fully connected hub with only one promoter and N enhancers would contain N enhancer-promoter interactions and (N)! / 2(N-2)! enhancer-enhancer interactions. For all values greater than N = 3, there would be more enhancer-enhancer loops than enhancer-promoter loops. To determine if our loops were forming such hubs we built interaction networks and detected communities of interacting anchor regions using a fast greedy modularity optimization algorithm (Clauset et al., 2004). We then classified these communities into two subsets: those containing a gained loop and those without. Twenty randomly chosen communities representing each subset are shown in figure 4F-G. Inspection of these subsets revealed stark differences. First, communities involving gained loops contained significantly more loop anchors (mean = 8.3 vs 3.6) compared to communities lacking gained loops (Wilcoxon Rank Sum Test, p = 3.7 * 10^−32^; Fig. 4H). This finding holds true even when accounting for the higher likelihood of a gained loop falling within a larger community (permutation testing, P < 0.001; Fig. S3). Second, the average ratio of enhancers to promoters per community was significantly higher (mean = 3.4 vs 1.6) in communities containing a PMA-specific loop (Wilcoxon Rank Sum Test, p = 7.4 * 10^−15^; Fig. 4I). Since communities involving gained loops had, on average, more than 3 enhancers it is consistent to observe more enhancer-enhancer than enhancer-promoter interactions in this subset.

These findings demonstrate that loops gained and activated during differentiation form activation hubs linking multiple distal enhancers to gene promoters (Kim and Dean, 2012; Krivega and Dean, 2016). Such interactive hubs have been previously reported including one that forms during erythroid differentiation bringing multiple distal DNase hypersensitive sites into close proximity with either fetal or adult globin genes (Palstra et al., 2003; Tolhuis et al., 2002). However, to the best of our knowledge this is the first report that such hubs are the primary location of newly formed and activated loops during cellular differentiation. Whether this phenomenon is specific to macrophage development or a more broadly applicable aspect of cellular differentiation remains to be seen.

### Dynamic re-wiring of chromatin architecture links distal AP-1 binding sites to key macrophage regulatory genes

Active regulatory elements can be profiled using assays that exploit chromatin accessibility to identify open chromatin regions (Boyle et al., 2008; Buenrostro et al., 2015; Giresi et al., 2007). To better characterize the cell-stage specific loops observed in our high resolution in situ Hi-C experiments, we mapped transposase-accessible DNA genome-wide by ATAC-seq in THP-1 cells. Comparison of chromatin accessibility before and after treatment uncovers thousands of changes in accessible chromatin regions during macrophage development. Analysis of the exact transposase insertion sites at each region of accessibility reveals regions of the genome that are protected from transposase activity, often representing genomic elements bound by transcription factors. Analysis of the sequence underlying such “footprints” can be used to predict which protein(s) may be present using an approach called TF footprinting (Fig. S4A). Global transcription factor footprinting of deeply sequenced ATAC-seq libraries in THP-1 cells both before and after PMA treatment reveals significant upregulation of protein-DNA interactions at JUN, FOS, and other AP-1-related target sequences, and down-regulation at sites targeted by interferon regulatory factors IRF8 and IRF9 (Fig. S4B). In agreement with these results, analysis of differential TF transcript abundance before and after PMA treatment reveals significant upregulation of genes encoding AP-1 family transcription factors, particularly for specific members of the MAF, JUN, and FOS AP-1 subfamilies, and significant downregulation of IRF8 (Fig. S4C). The upregulation of AP-1 transcription factor levels and corresponding increase in TF occupancy at AP-1 target sequences is consistent with the broad role of AP-1 in controlling cellular differentiation and proliferation (Eferl and Wagner, 2003; Shaulian and Karin, 2002).

We next leveraged TF footprinting analysis to specifically identify the transcription factors present at anchor regions of each class of loops (Fig. 5A-E). As expected, CTCF binding is strongly enriched at static chromatin loops, consistent with both motif enrichment and ChIP-seq experiments in THP-1 cells (Fig. 5C). Additional TF footprints, such as NRF1 and EGR1, are also statistically enriched but present at a small subset of static long-range interactions. Similar levels of CTCF are observed at lost and deactivated loop anchors. Analysis of gained and activated looping events, on the other hand, reveals a striking enrichment for AP-1 target sequences at more than 40% of loop anchors (Fig. 5D-E). Interestingly, CTCF footprints are far less enriched at gained and activated interaction sites. A careful inspection of ChIP-seq data reveals that CTCF is bound at these loop anchors but that CTCF binding is significantly weaker compared to static loop anchors. The functional significance of this difference is unclear but it further underscores the characteristic differences between static and gained/activated loops (Fig. S5A-C).

AP-1 binding was particularly enriched at gained and activated loop anchors with active enhancers (Fig. 5F), suggesting that AP-1 may be positively regulating gene transcription *via* binding at distal regulatory elements. To explore this possibility, we determined the relationship between distal TF binding and gene expression (Fig. 5G). Indeed, distal binding of AP-1 proteins, as measured by TF footprinting, correlated with increased expression of connected genes (Fig. 5G). While increased expression of genes that looped to distal FOSJUN motifs was particularly prominent, other FOS, JUN, and MAF family protein footprints showed a similar effect.

**Figure 5:**
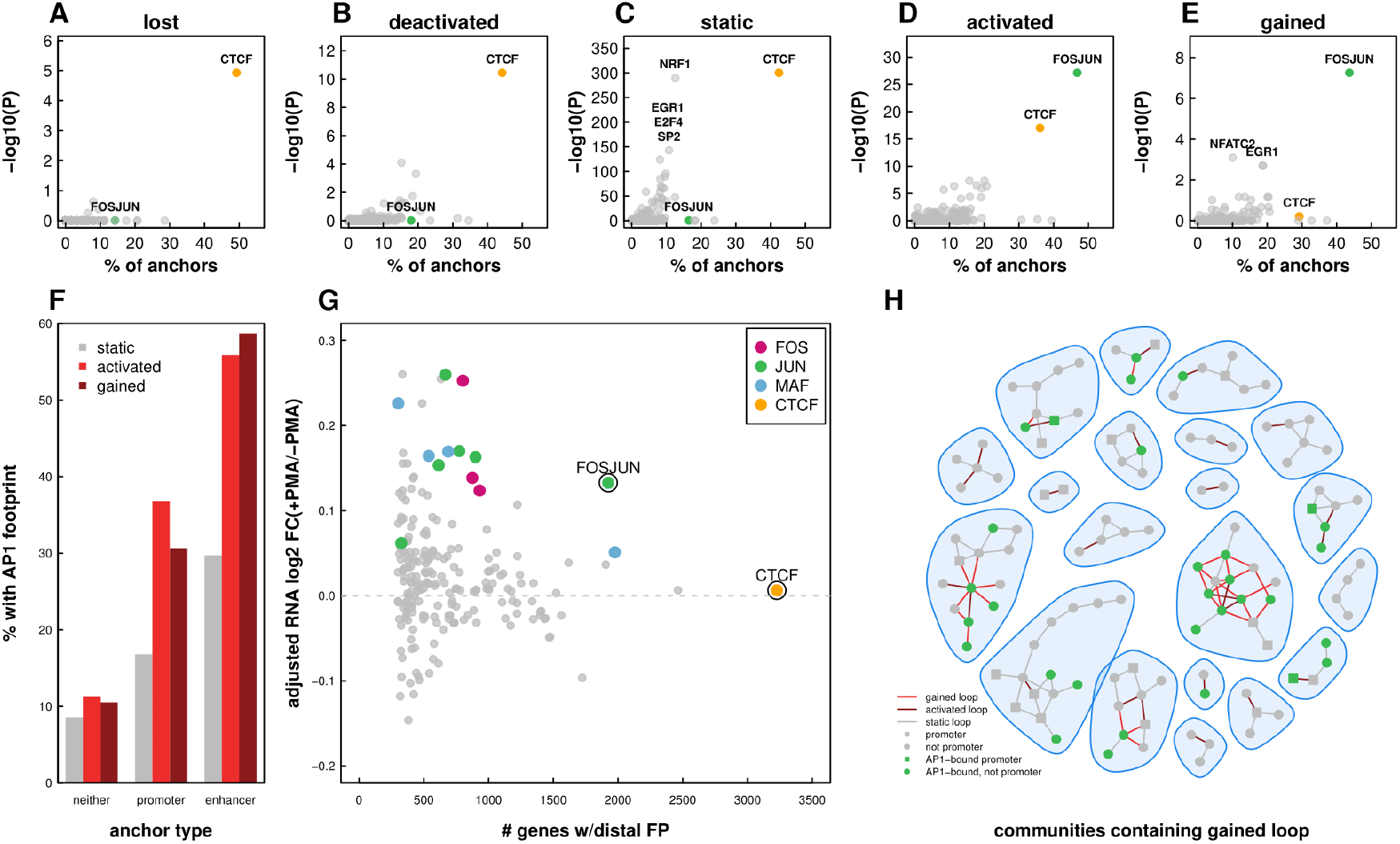
AP-1 enriched at enhancer containing loop anchors in both gained and activated loops. Scatter plots depicting the percent of lost (**A**), deactivated (**B**), static (**C**), activated (**D**), and gained (**E**) anchors that overlap TF footprints and the −log 10 p-value of enrichment (Fisher’s Exact Test) for each TF. (**F**) Bar plots depicting the percent of loop anchors that overlap an AP-1 footprint as a function of loop subset and promoter or enhancer overlap (**G**) Median log2 RNA fold change of genes connected via a loop to distal TF footprints. Log 2 FC were median normalized to account for the fact that genes at loop anchors exhibited a shift towards upregulation during differentiation. (**H**) 20 randomly chosen interaction communities containing a gained loop are shown. Loop anchors containing AP-1 footprints are indicated in green. See also Figure S4.

Taken together, the results presented here reveal multiple characteristics of promoter-distal gene regulation during macrophage development. Enhancer activation and AP-1 binding at distal ends of both gained and pre-formed loops create multi-loop activation hubs. These hubs harbor more that 3 distal regulatory elements per promoter and are associated with large increases in gene transcription. A clear example of a newly formed activation hub that exhibits all of these characteristics is observed at the TPRG1/BCL6 locus on chromosome 3 (Fig. 6). A complex network of AP-1 bound loci, located primarily within introns of the *LPP* gene, connect distal enhancers to the promoter regions of *TPRG1* and *BCL6*. Intriguingly, *LPP*, whose promoter is not involved in any gained or activated loops, exhibits only a moderate change in gene expression (1.4-fold increase). In contrast, *TPRG1* and *BCL6*, which are both associated with gained and activated loops, increase in expression by more than 25 fold.

**Figure 6:**
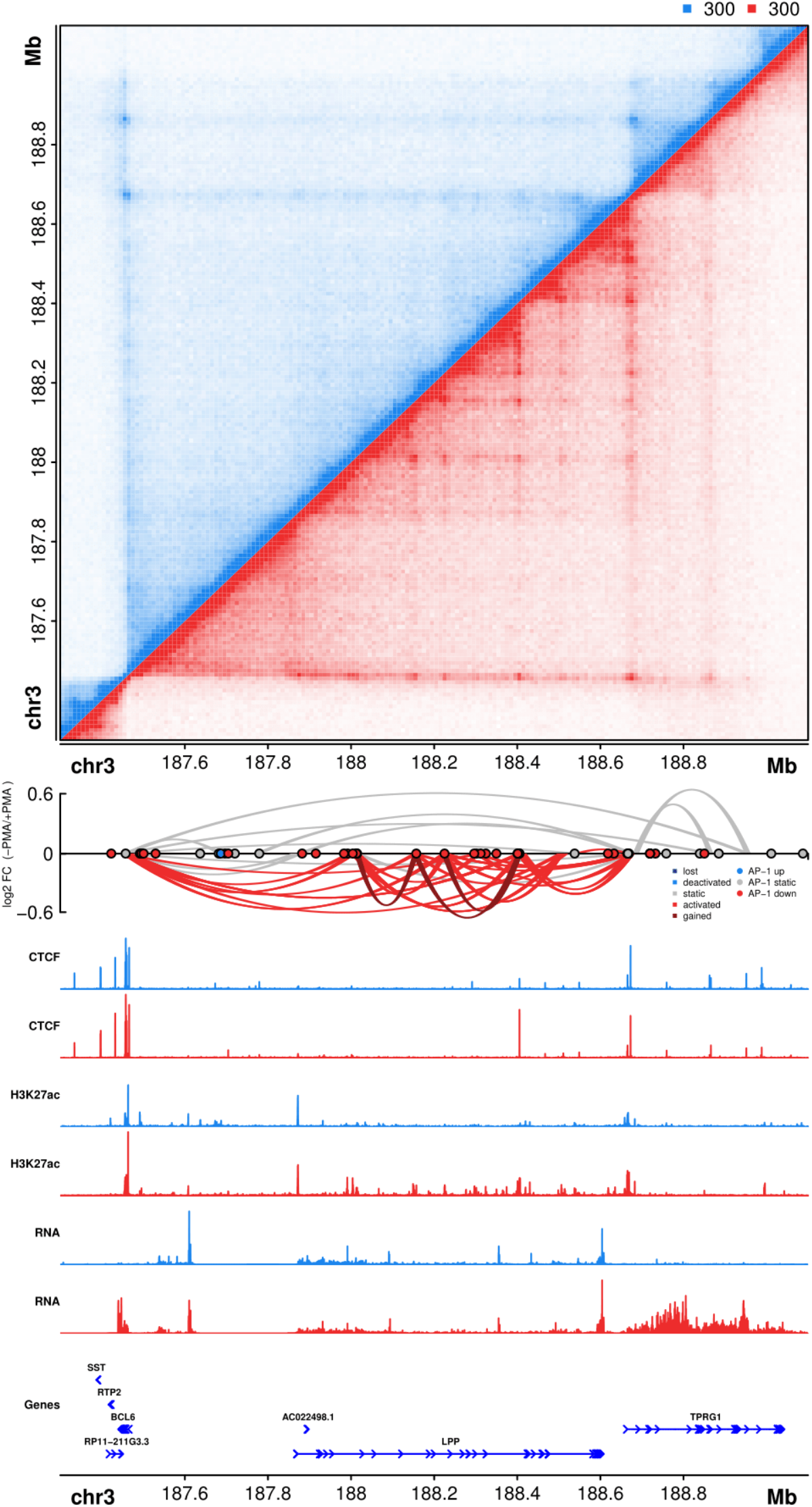
AP-1 bound activation hub formed during PMA induced differentiation of THP-1 cells on chromosome 3. (Top) Hi-C contact matrix depicting normalized contact frequencies between loci on a region of chromosome 3 in untreated THP-1 cells (blue, top left) and PMA treated THP-1 cells (red, bottom right). (**Middle**) A Sushi ribbon plot depicting DNA loops, loop fold changes (y-axis), loop subset (color of loops), and differential AP-1 footprints (circles). (**Bottom**) ChlP-seq signal tracks, RNA-seq signal tracks, and gene structures. See also Figures S5 and S6.

Further examples of AP-1 bound activation hubs are observed at key macrophage regulatory genes including *MAFB*, and *IL1β* (Fig. S6A-C). Transcriptional activation of *MAFB*, for example, is accompanied by dramatic changes in surrounding chromatin interactions, particularly for interactions between the *MAFB* gene promoter and a distal AP-1-bound enhancer that gains chromatin accessibility and TF occupancy in THP-1 derived macrophages (Fig. S6A). Activation of the interleukin-1b gene similarly involves dynamic looping between the *IL1β* gene promoter and a distal regulatory element marked by AP-1 binding after differentiation (Fig. S6B). The gain of AP-1 enhancer-promoter interactions are in certain cases marked as well by CTCF binding, suggesting AP-1 binding may be directed towards gene promoters for gene activation by CTCF (Fig. S6A). However, in other cases dynamic AP-1 looping-interaction sites are not marked by CTCF binding (Fig. S6C), or marked by nondynamic CTCF binding (Fig. S6B), suggesting the chromatin interactome at these genes can be directed either through AP-1-mediated interactions or through additional factors.

## DISCUSSION

Our results provide a new framework for understanding how the three-dimensional arrangement of human chromosomes may contribute to cellular differentiation and cell type-specific expression patterns. By generating deeply sequenced in situ Hi-C experiments in the context of monocyte-to-macrophage differentiation, we present some of the highest quality genome-wide maps of chromatin interactions during cellular differentiation to date. Identification of DNA loops in monocytes and THP-1 derived macrophages revealed multiple classes of preformed and dynamic 3D chromatin structures that correlate with changes in enhancer activation and gene expression. Specifically, we found that during macrophage development AP-1 binds at promoter-distal enhancer elements that are connected to upregulated gene promoters via a complex network of both gained and preexisting DNA loops. The multi-interaction AP-1 communities we identified contained multiple enhancers per promoter and were enriched for macrophage-related genes, suggesting that sophisticated 3D interaction networks may play an important role in developmental gene regulation. The AP-1 signature observed at activated and gained macrophage loops further suggest that transcription factor identity, which may be cell-type specific, may play an important role in driving dynamic transcription of target genes.

AP-1, a heterodimeric transcription factor comprising various combinations of FOS, JUN, MAF, ATF, and CREB family proteins, has been known to play a pivotal role in leukocyte development for decades (Liebermann et al., 1998; Valledor et al., 1998). However, its participation in gene regulation via DNA looping during macrophage development has not been previously described. Nevertheless, locus- and gene-specific examples of AP-1 bound DNA loops have been reported, such as at regulatory enhancer-promoter interactions that drive transcription of *ZEB2* and *PADI3* (Chavanas et al., 2008; Qiao et al., 2015), supporting a role for AP-1 family proteins in three-dimensional regulation of target genes and the broader, genome-wide participation of AP-1 characterized in the present study. Given its role across diverse cellular differentiation pathways (Eferl and Wagner, 2003; Shaulian and Karin, 2002), we speculate that the heterodimeric composition of the AP-1 transcription factor complex may contribute to re-wiring of chromatin interactions in a cell-type and tissue-specific manner. However, given the extraordinary number of potential transcription factor combinations that may co-bind at AP-1 consensus motifs (Mechta-Grigoriou et al., 2001), which can not be determined directly by footprinting methods, future studies aimed at comprehensively mapping this combinatorial landscape would shed significant insight into the precise proteins underlying AP-1 related looping events.

The upregulation of macrophage-related genes through both pre-existing DNA loops and through dynamic long-range interactions agrees with previous gene-specific examples of loop-dependent gene regulation within distinct developmental contexts. At the beta globin locus for instance, one of the best studied examples of long-range gene regulation (Kim and Dean, 2012), novel loop formation between locus control elements during blood cell development are required and sufficient for appropriate gene activation (Deng et al., 2012; Deng et al., 2014). In contrast, stimulation of IMR90 cells with TNFα activates enhancers at the promoter-distal anchors of pre-existing loops but does not induce large scale changes to 3D chromatin architecture (Jin et al., 2013). The identification of both static and dynamic loop-based mechanisms in various biological contexts suggests that both phenomena represent important paradigms for dynamic gene regulation. Moreover, our high quality maps of chromatin interactions in differentiating THP-1 monocytes reveals widespread interplay of static and dynamic looping events. In particular, we found that these two types of regulatory looping mechanisms co-occur at specific loci forming multi-loop activation hubs at key macrophage regulatory genes including *IL1β* and *MAFB*. These hubs often involve multiple distal enhancers looping to a single gene promoter and are associated with strong upregulation of gene transcription.

AP-1 bound activation hubs are reminiscent of dynamic chromatin structures found at the beta-globin locus in which multiple distal sites loop to the active beta globin genes during specific stages in erythroid cell development (Tolhuis et al., 2002). Analogous to AP-1 interactions targeting macrophage-specific genes, the beta-globin locus is organized into an “Active Chromatin Hub” that requires regulatory transcription factors such as ELKF, GATA1, Ldb1, and FOG1 (Drissen et al., 2004; Song et al., 2007; Vakoc et al., 2005). Indeed, we believe that the multi-interaction communities identified herein may have far-reaching implications for how chromosome organization instructs transcription in other cellular contexts and throughout human development. Identification of AP-1 activation hubs in our system was made achievable thanks to the high quality comprehensive interaction maps generated by deeply sequenced in situ Hi-C experiments. Thus, deep profiling of chromatin loops throughout early development remains a pressing need for better understanding the dynamic nature of mammalian chromosome organization.

Finally, because Hi-C and other chromatin profiling assays query DNA loops and DNA-protein interactions across cell populations, it is impossible to determine from these data sets whether all loops in a hub exist at the same time or if we are observing multiple subpopulations of cells with exclusive subsets of DNA looping events. Single cell profiling methods, while useful for comparison against aggregate cell populations, also struggle to discriminate between these two possibilities, as only a single anchor-to-anchor ligation event is generated per allele by chromosome conformation capture. Nevertheless, it is difficult to imagine a scenario in which AP-1 binding to one enhancer would preclude its binding, and perhaps consequently looping, to additional local regulatory elements. In either case, it is likely that activation hubs increase the local concentrations of enhancers and distally bound transcription factors at gene promoters contributing to increased transcription. Further development of computational methods and experimental methods to identify multi-loop communities, such as Concatemer Ligation Assay (“COLA”), should help address some of these pressing questions in chromatin structure and gene regulation (Darrow et al., 2016).

## EXPERIMENTAL PROCEDURES

### THP-1 cell culture and differentiation

THP-1 cells were obtained from ATCC and passaged in growth medium containing RPMI-1640 (Corning), 10% fetal bovine serum, and 1% penicillin streptomycin. THP-1 differentiation was carried out at a final concentration of 100 nM PMA for 72 hours, followed by trypsinization and isolation of adherent THP-1 derived macrophages with TrypLE (ThermoFisher).

### Chromatin Immunoprecipitation

For each biological replicate, approximately 10 million treated or untreated THP-1 cells were resuspended in growth media at 1 × 10^6^ cells/mL. Fixation was performed for 10 minutes with rotation at room temperature in 1% formaldehyde; cross-linking was then quenched in 200 mM glycine with rotation for 5 minutes. Cross-linked cells were then pelleted and resuspended in 1× RIPA lysis buffer, followed by chromatin shearing via sonication (3 cycles using a Branson sonicator: 30 seconds on, 60 seconds off; 15 additional cycles on a Bioruptor sonicator: 30 seconds on, 30 seconds off). Individual ChIP experiments were performed on pre-cleared chromatin using antibody-coupled Dynabead protein G (ThermoFisher) magnetic beads. Anti-histone H3 (acetyl K27) antibody was obtained from Abcam (ab4729), CTCF antibody was obtained from Millipore (07–729). 3–5 ug of antibody per ChIP was coupled to 18 uL beads and rotated overnight with sheared chromatin at 4^°^ C. Beads were then washed 5× in ChIP wash buffer (Santa Cruz), 1× in TE, and chromatin eluted in TE + 1% SDS. Cross-linking was reversed by incubation at 65^0^ C overnight, followed by digestion of RNA (30 min RNase incubation at 37^0^C) and digestion of protein (30 min proteinase K incubation at 45^0^C). ChIP DNA was then purified on a minElute column (Qiagen), followed by DNA library preparation and size selection of 350–550 bp fragments via gel extraction (Qiagen).

### Assay for transposase accessible chromatin (ATAC-seq)

For each biological replicate, approximately 50,000 treated or untreated THP-1 cells were collected and washed with 1x ice cold PBS. Cells were pelleted via centrifugation and resuspended in lysis buffer (10mM Tris-HCl, pH 7.4, 10mM NaCl, 3mM MgCl_2_⅛, 0.1% IGEPAL CA-630). Cells were again pelleted by centrifugation and the supernatant discarded. Transposition was carried out for 30 minutes at 37^0^C using the Nextera DNA library prep kit (Illumina, cat#FC-121-1030). DNA was subsequently purified on a minElute column (Qiagen), and PCR amplified using the NEBNext high-fidelity master mix (NEB cat#M0541) with nextera PCR primers and barcodes. PCR amplification was monitored as described (Buenrostro et al. 2015), and gel purified to remove contaminating primer-dimer species.

### In situ Hi-C

In situ as performed exactly as described by Rao et al (Rao et al., 2014). Cells were crosslinked in 1% v/v formaldehyde for ten minutes with stirring and quenched by adding 2.5M glycine to a final concentration of 0.2M for 5 minutes with rocking. Cells were pelleted by spinning at 300 G for 5 minutes at 4 degrees C. Cells were washed with cold PBS and spun again prior to freezing in liquid nitrogen.

Cells were lysed with 10mM Tris-HCl pH8.0, 10mM NaCl, 0.2% Igepal CA630 and protease inhibitors (Sigma, P8340) for 15 minutes on ice. Cells were pelleted and washed once more using the same buffer. Pellets were resuspended in 50ul of 0.5% SDS and incubates for 5–10 minutes at 62’C. Next reactions were quenched with 145ul of water and 25ul of 10% Triton X-100 (Sigma, 93443) at 37’C for 15 minutes. Chromatin was digested overnight with 25ul of 10X NEBuffer2 and 10U of MboI at 37’C with rotation.

Reactions were incubated at 62’C for 20 minutes to inactivate MboI and then cooled to room temperature. Fragment overhangs were repaired by adding 37.5 ul of 0.4mM biotin-14-dATP, 1.5 ul of 10mM dCTP, 1.5 ul of 10mM dGTP, 1.5 ul of 10mM dTTP, and 8 ul of 5U/ul DNA Polymerase I, Lar (Klenow) Fragment and incubating at 37’C for 1 hour. Ligation was performed by adding 667 ul of water, 120 ul of 10X NEB T4 DNA ligase buffer, 100 ul of 10% Triton X-100, 12 ul of 10 mg/ml BSA, and 1 ul of 2000 U/ul T4 DNA Ligase and incubating at room temperature for 4 hours. Samples were pelleted at 2500 G and resuspended in 432 ul water, 18 ul 20 mg/ml proteinase K, 50 ul 10% SDS, 46 ul 5M NaCl and incubated for 30 minutes at 55’C. The temperature was raised to 68’C and incubated overnight.

Samples were cooled to room temperature. 874 ul of pure ethanol and 55 ul of 3M sodium acetate, pH 5.2 were added to each tube which were subsequently incubated for 15 minutes at −80’C. Tubes were spun at max speed at 2’C for 15 minutes and washed twice with 70% ethanol. The resulting pellet was resuspended in 130 ul of 10mM Tric-HCl, pH8 and incubated at 37’C for 15 minutes. DNA was sheared using an LE220 Covaris Focused-ultrasonicator to a fragment size of 300–500 bp. Sheared DNA was size selected using AMPure XP beads. 110 ul of beads were added to each reaction and incubated for 5 minutes. Using a magnetic stand supernatant was removed and added to a fresh tube. 30ul of fresh AMPure XP beads were added and incubated for 5 minutes. Beads were separated on a magnet and washed two times with 700 ul of 70% ethanol without mixing. Beads were left to dry and then sample was eluted using 300 ul of 10 mM Tris-HCl, pH 8.

150 of 10 mg/ml Dynabeads MyOne Streptavidin T1 beads were washed resuspended in 300 ul of 10 mM Tris HCl, pH 7.5. This solution was added to the samples and incubated for 15 minutes at room temperature. Beads were washed twice with 600ul Tween Washing Buffer (TWB; 250 ul Tris-HCl, pH 7.5, 50 ul 0.5 M EDTA, 10 ml 5M NaCl, 25 ul Tween 20, and 39.675 ml water) at 55’C for 2 minutes with shaking. Sheared ends were repaired by adding 88 ul 1X NEB T4 DNA ligase buffer with 1mM ATP, 2 ul of 25 mM dNTP mix, 5 ul of 10U/ul NEB T4 PNK, 4ul of 3U/ul NEB T4 DNA polymerase I, 1ul of 5U/ul NEB DNA polymerase I, Large (Klenow) Fragment and incubating at room temperature for 30 minutes. Beads were washed two more times with TWB for 2 minutes at 55’C with shaking. Beads were washed once with 100 ul 1X NEBuffer 2 and resuspended in 90 ul of 1X NEBuffer 2, 5 ul of 10 mM dATP, 5ul of 5U/ul NEB Klenow exo minus, and incubated at 37’C for 30 minutes. Beads were washed two more times with TWB for 2 minutes at 55’C with shaking. Beads were washed once in 50 ul 1X Quick Ligation reaction buffer and resuspended in 50 ul 1X Quick Ligation reaction buffer. 2 ul of NEB DNA Quick ligase and 3 ul of an illumina indexed adapter were added and the solution was incubated for 15 minutes at room temperature. Beads were reclaimed using the magnet and washed two more times with TWB for 2 minutes at 55’C with shaking. Beads were washed once in 100 ul 10 mM Tris-HCl, pH8 and resuspended in 50 ul 10 mM Tris-HCl, pH8.

Hi-C libraries were amplified for 7–12 cycles in 5 ul PCR primer cocktail, 20 ul of Enhanced PCR mix, and 25 ul of DNA on beads. The PCR settings included 3 minutes of 95’C followed by 7–12 cycles of 20 seconds at 98’C, 15 seconds at 60’C, and 30 seconds at 72’C. Samples were then held at 72’C for 5 minutes before lowering to 4’C until samples were collected. Amplified samples were brought to 250 ul with 10 mM Tris-HCl, pH 8. Samples were separated on a magnet and supernatant was transferred to a new tube. 175 ul of AMPure XP beads were added to each sample and incubated for 5 minutes. Beads were separated on a magnet and washed once with 700 ul of 70% ethanol. Supernatant was discarded. 100 ul of 10 mM Tris-HCl and 70 ul of fresh AMPure XP beads were added and the solution was incubated for 5 minutes at room temperature. Beads were separated with a magnet and washed twice with 700 ul 70% ethanol. Beads were left to dry until cracking started to be observed and eluted in 25 ul of Tris HCl, pH 8.0. The resulting libraries were quantified by Qubit and Bioanalyzer prior to sequencing.

### RNA sequencing

For each replicate, approximately 5 million treated or untreated THP-1 cells were collected and washed with 1x ice cold PBS. RNA was extracted using the Dynabeads mRNA Direct kit according to manufacturer directions. Sequencing libraries were prepared using the Epicenter ScriptSeq V2 kit according to manufacturer supplied protocol. The resulting libraries were quantified by Qubit, mixed in equal concentrations, and assayed by Bioanalyzer prior to sequencing using Illumina HiSeq 2500 technology.

### Genomic data processing

#### ATAC-seq

Paired end sequencing reads were trimmed using trim galore version 0.40 with command-line settings “trim_galore -q 20 --trim1 --paired” and subsequently aligned to the hg19 reference genome using bowtie2 version 2.2.4 with settings “bowtie2 -t --sensitive”. Mapped reads were merged across technical and sequencing replicates, and duplicate reads removed using picard tools version 1.92. Peaks were identified for each sample and biological replicate using MACS2 version 2.1.0 with command line options “macs2 callpeak --bdg --nomodel -t -g hs”. For analysis of differential chromatin accessibility, raw ATAC-seq reads were extracted for each condition over a merged set of ATAC-seq peaks and statistical significance determined using the glmTreat function in edgeR with a fold change cutoff of 2. For genome-track visualization purposes, ATAC-seq reads were normalized for sequencing depth and mappability using the align2rawsignal pipeline with options “-of=bg -n=5 -l=200 -w=200 -mm=30” (<u>https://align2rawsignal.googlecode.com</u>).

#### ChIP-seq

Paired end sequencing reads were trimmed using trim galore version 0.40 with command-line settings “trim_galore -q 20 --trim1 --paired” and subsequently aligned to the hg19 reference genome using bowtie1 version 1.1.1 with settings “bowtie -q --phred33-quals -X 2000 -m 1 --fr -p 8 -S -- chunkmbs 400”. Mapped reads were merged across technical and sequencing replicates, and duplicate reads removed using picard tools version 1.92. Peaks were identified for each sample and biological replicate using MACS2 version 2.1.0 with command line options “macs2 callpeak -- bdg -t -g hs”. For analysis of differential chromatin accessibility, raw ATAC-seq reads were extracted for each condition over a merged set of ATAC-seq peaks and statistical significance determined using the glmTreat function in edgeR with a fold change cutoff of 2. For genome-track visualization purposes, ATAC-seq reads were normalized for sequencing depth and mappability using the align2rawsignal pipeline with options “-of=bg -n=5 -l=200 -w=200 -mm=30” (<u>https://align2rawsignal.googlecode.com</u>).

#### RNA-seq

Paired end sequencing reads were trimmed using trim galore version 0.40 with command-line settings “trim_galore -q 20 --trim1 --paired” and subsequently aligned to gencode v19 transcripts using kallisto with default parameters. For analysis of differential genes, kallisto outputs were processed using tximport. Genes with less than 2 counts per million were filtered out. Statistical significance was determined using the glmTreat function in edgeR with a fold change cutoff of 2. For genome-track visualization purposes, RNA-seq reads were aligned to the hg19 reference genome using tophat. Duplicate reads were removed using MarkDuplicates from picard-tools version 1.92. The resulting reads were normalized for sequencing depth and mappability using the align2rawsignal pipeline with options “-of=bg -n=5 -l=1 -w=200 -mm=30” (<u>https://align2rawsignal.googlecode.com</u>).

#### Hi-C

In situ Hi-C data sets were processed as described by Rao et al with minor differences to FDR calculations and final filtering parameters. Mbol fragments of the human hg19 reference genome were determined using hicup_digester with the following command: “hicup_digester -g Human_hg19 -re1 ^GATC,Mbol”. Hi-C fastq files were split into small files of 5 million reads each. Reads were aligned to the human reference genome hg19 using bwa mem version 0.7.12 with default parameters. Read pairs, in which both ends mapped uniquely were retained. Reads likely to mapping of a ligation junction were also retained as described by Rao et al (Rao et al., 2014). Each read was assigned to a single fragment using bedtools. Read pairs were filtered for unique combinations of chromosomes, start positions, and strand orientations to remove potential artifacts from PCR duplication. Such filtering was applied after merging sequencing duplicates but prior to merging data sets arising from different library preparations. Filtered reads from each pair of SAM files were combined into a single bed paired end file. Reads with mapping scores (MAPQ) below 30 were filtered out from subsequent analyses. We next built contact matrices for various resolutions including 5, 10 and 100Kb. For each resolution we binned all fragments according to their midpoint and then counted the read pairs that corresponded to each pair of fragments. Only intra chromosomal matrices were constructed. We constructed two type of contact matrices. For differential loop calling we built matrices for each biological replicate separately. For visualization we combined biological replicates into a single contact matrix for each sample (i.e. un-treated and PMA-treated THP-1 cells). Matrices were balanced according to a method proposed by Knight and Ruiz (Knight and Ruiz, 2013). Bins with less than 25 pixels of non-zero values were discarded from the normalization procedure. After balancing we calculated the expected normalized contacts for each distance for each chromosome separately. Noise associated with distances with few counts was mitigated by merging distances until more than 400 counts were achieved.

P values describing the observed contact frequencies given local background contact frequencies were determined for pixels at 10Kb resolution as described by Rao et al. For all pixels representing genomic bins separated by less than 2 million base pairs, various metrics were collected. For each pixel we defined several local neighborhoods as described by Rao et al; donut, horizontal, vertical, and lower right. Values of p=2 and w = 5 were used. The donut neighborhood is defined as pixels that are greater than p and less than or equal to w pixels away from the primary pixel in either the x or y directions. The other three neighborhoods are subsets of the donut neighborhood. The horizontal neighborhood is defined as pixels that are greater than p and less than or equal to w pixels away from the primary pixel in the x direction and greater less than p pixels away in the y direction. The horizontal neighborhood is defined as pixels that are greater than p and less than or equal to w pixels away from the primary pixel in the y direction and less than p pixels away in the x direction. And the lower right neighborhood is all pixels with x values greater than the primary pixel but less than or equal to w pixels away, pixels with y values less than than the primary pixel and less than or equal to w pixels away, and pixels with coordinates that are more than p pixels away in either the x or y direction. For each neighborhood, summed normalized contact frequencies were determined. If summed values were less than 16, w was increased until either greater than 16 counts were reached or w was equal to 20. For each neighborhood, summed expected contact frequencies were determined as well. For each neighborhood the ratio of observed / expected counts was determined. This ratio was multiplied by the expected value of the primary pixel to determine the expected normalized contacts for each pixel analyzed. This value was converted to an expected raw contact count by multiplying by the corresponding normalization factors determined by our matrix balancing step. P values of differences between observed raw counts and expected raw count, as estimated from each of the four local neighborhoods, were was determined determined using the R programming language and the function ppois with lower.tail = FALSE. P values for all pixels tested on all chromosomes were corrected for multiple hypothesis testing using the Benjamini-Hochberg method.

Loops were determined by further clustering and filtering of significant pixels in a similar fashion as described by Rao et al. All pixels with corrected p values less than or equal to 0.05 were clustered using the DBSCAN algorithm with epsilon = 20 Kb. For each cluster the pixel with the highest normalized counts was retained and annotated with the number of pixels in its cluster. These pixels were filtered for the following parameters; fold-change of observed vs expected (donut) > 1.75, fold-change of observed vs expected (horizontal) > 1.5, fold-change of observed vs expected (vertical) > 1.5, fold-change of observed vs expected (lower right) > 1.5, the sum of the adjusted p-values for all four neighborhoods < 0.001, and the number of pixels in the cluster > 2. Finally, to avoid artifacts due to local neighborhoods that contained regions of repetitive nature, pixels within 50Kb of a genomic bin that was discarded by the matrix normalization step were removed.

#### TF Footprinting

Transcription factor footprinting was broken into two steps: (A) identify bound TF motifs in THP-1 monocytes and THP-1 derived macrophages and (B) Determine differential footprinting scores before and after PMA treatment for each motif identified in THP-1 cells.

#### Step (A)

Putative TF footprints were identified using the Protein Interaction Quantification (PIQ) footprinting algorithm (Sherwood et al., 2014) against the JASPAR core vertebrate database of TF motifs (<u>http://jaspar.genereg.net</u>). First, motif matching was performed for 516 known TF target sequences against the hg19 reference genome using the PIQ package pwmmatch.exact.r script. Second, filtered ATAC-seq alignment reads were converted into binary RData files using the PIQ package pairedbam2rdata.r script. Third, TF footprint scores were determined for each motif match using the PIQ package pertf.bg.r and common.r scripts with default settings. Putative TF footprints were filtered at a positive predictive value (PPV) cutoff of 0.7 and for footprints that intersect ATAC-seq peaks identified in THP-1 cells. Altogether, 2,731,616 TF footprints were identified post-filtering with a median 3,693 binding sites per unique TF motif.

#### Step (B)

Analysis of dynamic TF binding was performed using the Wellington-bootstrap algorithm for differential footprinting (Piper et al., 2015) against all post-filtering TF footprints identified by PIQ. First, differential footprinting was applied using the pyDNase wellington-bootstrap.py script with the command-line option for ATAC-seq input “-A”. Second, differential footprint scores were determined using the pyDNase dnase_ddhs_scorer.py script with the command-line option for ATAC-seq input (“-A”). Differential footprint scores (DFP) were altogether adjusted by median normalization, followed by median differential footprint analysis for each independent factor. Wellington DFPs strongly correlate with changes in PIQ PPV values (p < 2.2e-16), and using a cutoff of +/− 2 standard deviations, we identify a total of 33,130 decreasing and 95,898 increasing TF binding events.

### Post processing and analyses

#### Comparison to Juicer pipeline

Hi-C libraries were also analyzed using the Juicer pipeline. Data was processed for 5,929,803,301 Hi-C read pairs in untreated THP-1 cells, yielding 3,935,374,088 Hi-C contacts and 5,522,487,839 Hi-C read pairs in PMA-treated THP-1 cells, yielding 3,789,121,851 Hi-C contacts. Loops were annotated using HiCCUPS at 5kB and 10kB resolutions with default Juicer parameters. This yielded a list of 14,964 loops in untreated THP-1 cells and 22,615 loops in PMA-treated THP-1 cells. All the code used in the above steps is publicly available at (github.com/theaidenlab). Loops identified by Juicer were filtered for those that were greater or equal to 50Kb and less than or equal to 2Mb. In order to determine overlap between our loops and Juicer we had to consider not only the pixel that was picked to represent the loop but all enriched pixels within a cluster. Therefore, for Juicer loops we picked the centroid of the cluster +/− the radius of the cluster for each loop anchor. We then rank ordered our loop calls by increasing p values and determined the percent of our loops that were also found in the Juicer loops calls. We did this for 20 subsets of our loop calls ranging from the top 5% of our loop calls to 100% of our loop calls. These overlaps were performed for both data sets.

#### Enhancer definition

For all of the analyses in this paper enhancers were defined as regions with an H3K27ac peak as determined by ChIP-seq. H3K27ac peaks that overlapped a gene promoter (i.e. the 2000 bp region upstream of a UCSC hg19 known gene transcription start site) were removed from this list.

#### Hi-C normalization for visualization

Visualizing differences between to Hi-C contact matrices can be complicated by different sequencing depths between data sets as well as differences in average interaction frequencies as a function of distance. To allow for accurate visual comparison of contact matrices between two samples all PMA-treated matrices were normalized such that the median normalized contact frequency for each genomic distance was identical between untreated and PMA-treated cells. For all distances plotted, the median contact frequency was determined for each data set. For each distance an offset was determined by dividing the median untreated value but the median PMA-treated value. All PMA-treated normalized contact frequencies were multiplied by this offset factor prior to plotting.

#### Aggregate peak analysis

In order to assess the quality of loop calls we generated aggregate peak analysis (APA) plots and scores. These analyses aggregate the signal of pixels of loops as well as the pixels surrounding them, the local background. For each loop in a given set of loops, the normalized observed contact frequencies were collected for the pixel representing the loop as well as for pixels within 10 bins in both the x and y directions. To normalize for loops at different distances, each pixel was divided by the expected normalized interaction frequency at that distance to give an observed over expected ratio. Median observed over expected ratios for each position in the matrix were calculated and plotted as a heatmap. APA scores were determined by dividing the center pixel value by the median value of the nine pixels in the lower right section of the APA plot.

#### Motif enrichment at loop anchors

To determine the frequency of transcription factor motif overlap with loop anchors, we downloaded TF motifs determined by Factorbook and intersected the motifs with our 10Kb loop anchors using the bedtools intersect function (Quinlan and Hall, 2010; Wang et al., 2013). To determine the expected frequency of overlap we shuffled the assignments of motif and genomic region 100 times and performed the same analysis. Unannotated motifs, those that started with “UAK” were not included.

#### CTCF motif orientation analysis

Loops were filtered to retain those that overlapped a CTCF binding site, as determined by ChIP-seq, at both anchors. These loops were further filtered for those that overlapped a single CTCF motif as determined by Factorbook (Wang et al., 2013). We then calculated the percent of those remaining loops that contained each of the four possible pairwise combinations of motifs.

#### Differential Loop analysis

Detection of differential loops can be complicated by larger scale changes chromatin structure that do indeed change interaction frequency but do not change looping per se. Such large-scale structural changes include changes in chromatin compaction, changes in domain boundaries, and duplication of genomic regions. To specifically detect changes in looping we devised a method that looked for changes in the enrichment of pixels representing DNA loops compared to local background interaction frequencies.

First, we collected information for all pixels that were identified as loops in either untreated or PMA-treated THP-1 cells. For each loop pixel we collected the following information from each sample and each biological replicate: raw interaction counts, expected interaction counts, and normalization factors. We collected that same information for local background pixels defined as pixels that were greater than 2 and less than 6 pixels away from the loop pixel in both x and y directions. Using this data we built a DESeq2 counts matrix and sample table as follows. Counts for all pixels in all samples in all biological replicates formed the columns of the count matrix and the rows represented different loops. A corresponding table described the relationship among the columns of the count matrix: each pixel belonging to a sample, a biological replicate, and either representing a loop ‘L’ or a background ‘B’ pixel. We analyzed this data for differential enrichment of the loop over background across condition, using the DESeq2 design formula “~ rep + sample + rep:sample + pixel_type + sample: pixel_type”. Raw counts required normalization to account for differential sequencing depth in each genomic bin and distance-dependent interaction frequencies, while sample-specific sequencing depth was controlled via terms in the design formula. A per-pixel normalization factor was calculated by multiplying the normalization factor for bin 1, the normalization factor for bin 2, and the expected interaction frequency. These normalization factors were centered around a value of 1 and added to the DESeq2 analysis. The DESeq2 differential pipeline was run with settings betaPrior=FALSE and dispersion fitType="local". The resulting loops were filtered for those with a p value of < 0.001 to produce our final set of differential loops.

#### Gene Ontology enrichment

GO enrichment was performed on genes whose promoter overlapped either a gained or activated loop. Promoters were defined as 2000 base pair regions upstream of a gene TSS. For activated loops, only genes at the distal end of an upregulated H3K27ac mark were considered. The background gene set used for these analyses was a list of all genes whose promoters overlapped a loop anchor in either treated or untreated cells. GO enrichment was performed for biological process gene ontologies using the goana function from the R package limma (Ritchie et al., 2015). GO terms were filtered for adjusted p value < 0.05. Selected enriched GO terms were plotted.

#### Loop community detection and analysis

Non-directed graphs were constructed from loops using the R package igraph. Communities were determined using the fastgreedy.community function with default parameters.

#### Footprint enrichment at loop anchors

To determine enrichment of TF at loop anchors we first intersected all TF footprints with loop anchors using the bedtools intersect function. For each TF we determined the number of footprints of that TF that overlapped an anchor, number of footprints of that TF that did not overlap an anchor, the number of footprints of other TFs that overlapped an anchor, and the number of footprints of other TFs that did not overlap an anchor. Using these values, we built a contingency table and performed fishers exact test in R. The resulting p values were corrected for multiple hypothesis testing using the Benjamini-Hochberg procedure.

## ACCESSION NUMBERS

The sequencing data from this study is publicly available through the Gene Expression Omnibus (GEO) series number GSE96800 (ChIP-seq, ATAC-seq, RNA-seq) and the Sequencing Read Archive (SRA) accession number PRJNA385337 (in situ Hi-C). Processed Hi-C data is also available through Juicebox (http://www.aidenlab.org/juicebox/).

## AUTHOR CONTRIBUTIONS

Conceptualization, DHP, KVP; Methodology, DHP, KVP, DVS; Software, DHP, KVP, IA, MSS, MIL, ELA; Formal Analysis, DHP, KVP; Funding Acquisition, DHP, MPS; Visualization, DHP, KVP; Writing, DHP, KVP; Validation, GTH, MCB.

## ACKNOWLEDGEMENTS

DHP is supported by National Institutes of Health, National Human Genome Research Institute (NHGRI) grant R00HG008662 and the Damon Runyon Cancer Research Foundation (DRG-2122-12). KVB is supported by National Institutes of Health, National Institute of Diabetes and Digestive and Kidney Diseases (NIDDK) grant F32DK107112. GTH is supported by NIH grant NIH T32HG000044. MIL is supported by National Institutes of Health, National Cancer Institute (NCI) grant CA142538-07. DVS was supported by an NIH T32 fellowship (HG000044) and a Genentech Graduate Fellowship. This research was supported by NIH Centers of Excellence in Genomic Science (CEGS) grant 5P50HG00773502. ELA is supported by an NIH New Innovator Award (1DP2OD008540), an NSF Physics Frontiers Center Award (PHY-1427654, Center for Theoretical Biological Physics), the National Human Genome Research Institute (NHGRI) Center for Excellence for Genomic Sciences (HG006193), the Welch Foundation (Q-1866), an NVIDIA Research Center Award, an IBM University Challenge Award, a Google Research Award, a Cancer Prevention Research Institute of Texas Scholar Award (R1304), a McNair Medical Institute Scholar Award, an NIH 4D Nucleome Grant U01HL130010, an NIH Encyclopedia of DNA Elements Mapping Center Award UM1HG009375, and the President’s Early Career Award in Science and Engineering. Illumina sequencing services were performed by the Stanford Center for Genomics and Personalized Medicine. The content is solely the responsibility of the authors and does not represent the official views of the National Institutes of Health. We would like to thank Craig D Wenger and Neva C Durand for helpful advice, guidance, and discussions.

---

**Figure S1:**
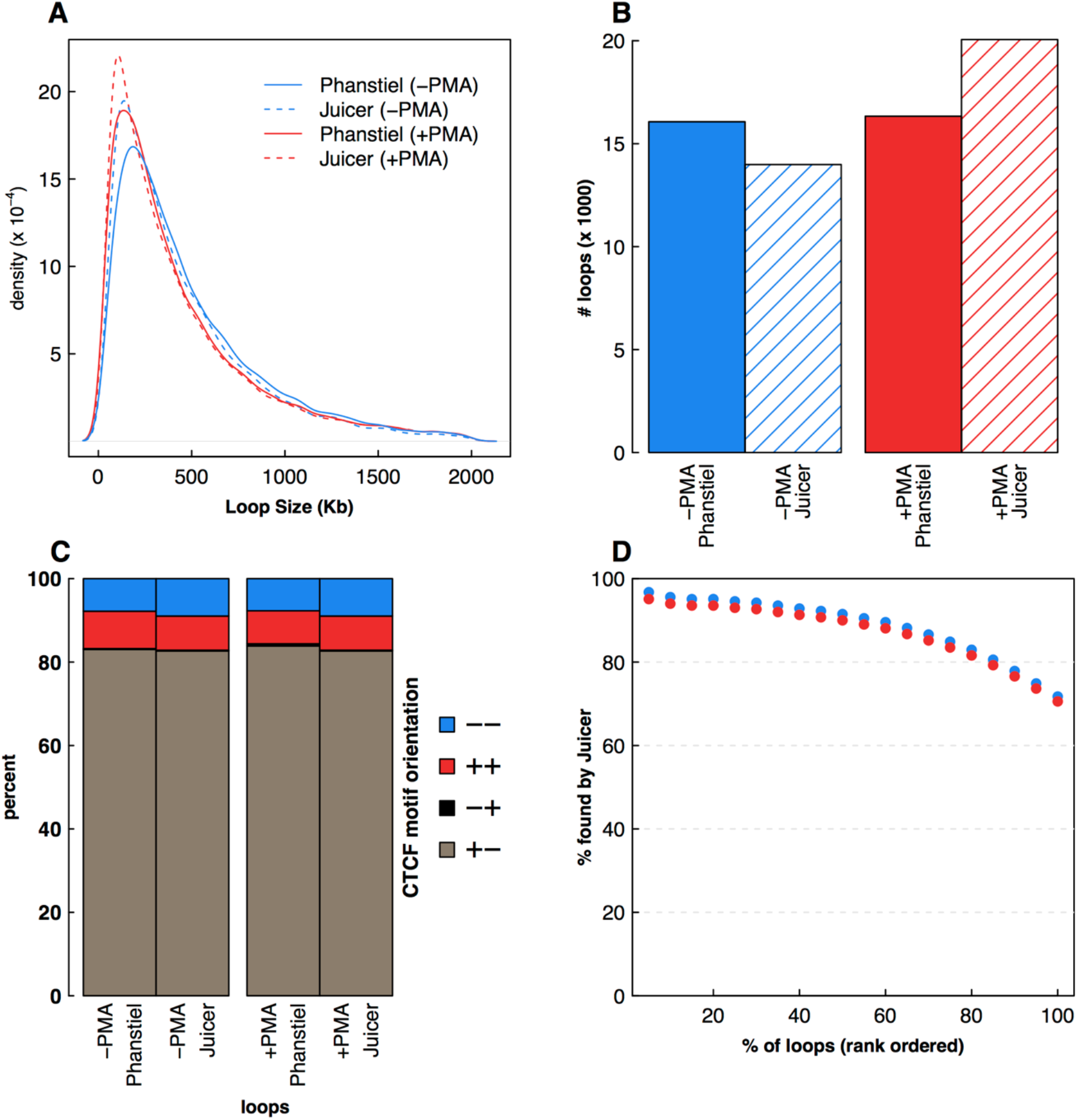
Comparison to loops detected by Juicer, Related to Figure 1. (**A**) A density plot depicts size distribution of loops detected by our method vs Juicer. (**B**) Barplots depict the number of loops detected by by our method vs Juicer for each data set. (**C**) Stacked bar plot depicting CTCF motif orientations at loop anchors as a percentage of all loops that contain a single CTCF bound peak at each anchor. (**D**) A scatter plot depicts the percent of loops that were also detected by Juicer as a function of rank order.

**Figure S2:**
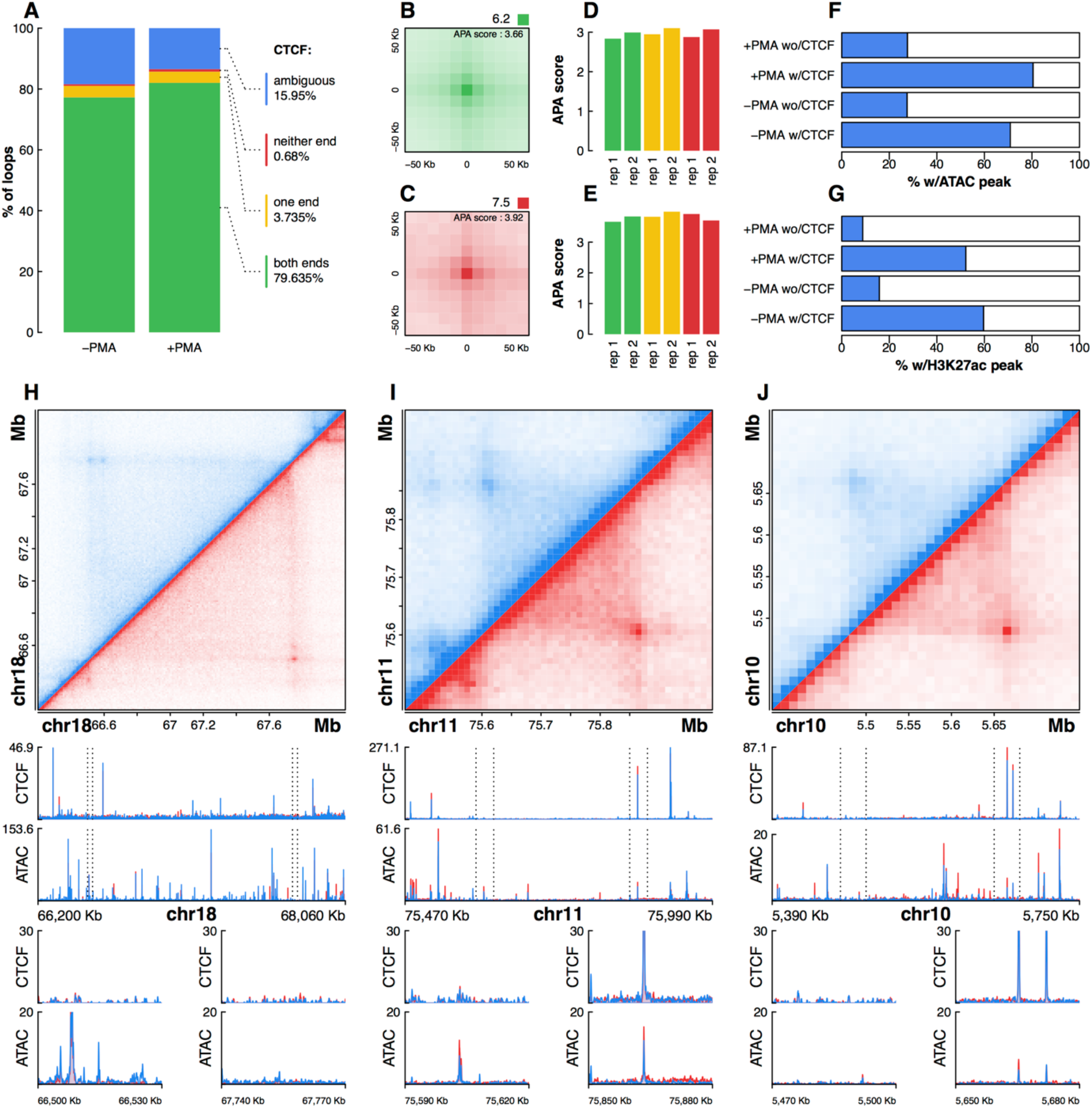
Non-CTCF-bound loops in untreated and PMA-treated THP-1 cells, Related to Figure 1. (**A**) A stacked bar plot depicts the percent of loops in each data sets that had a CTCF binding site, determined by ChIP-seq, at zero, one, or two ends. Loop anchors which lacked a CTCF binding site but were within 10Kb of a CTCF binding site were categorized as ‘ambiguous’. APA plots for loops with CTCF at both (**B**) or neither (**C**) end. Bar plots depicting APA scores for subsets of DNA loops that were bound at one (yellow), two (green), or neither (red) end for both untreated (**D**) and PMA-treated (**E**) THP-1 cells. (**F**) The percent of loop anchors that overlap an ATAC-seq peak was determined for anchors that did and did not overlap a CTCF binding site. (**G**) The percent of loop anchors that overlap an H3K27 acetylation peak was determined for anchors that did and did not overlap a CTCF binding site. (**H-J**) Hi-C contract matrices, ChIP-seq signal tracks, and ATAC-seq signal tracks for three regions harboring loops that lack CTCF binding sites at one or both anchors.

**Figure S3:**
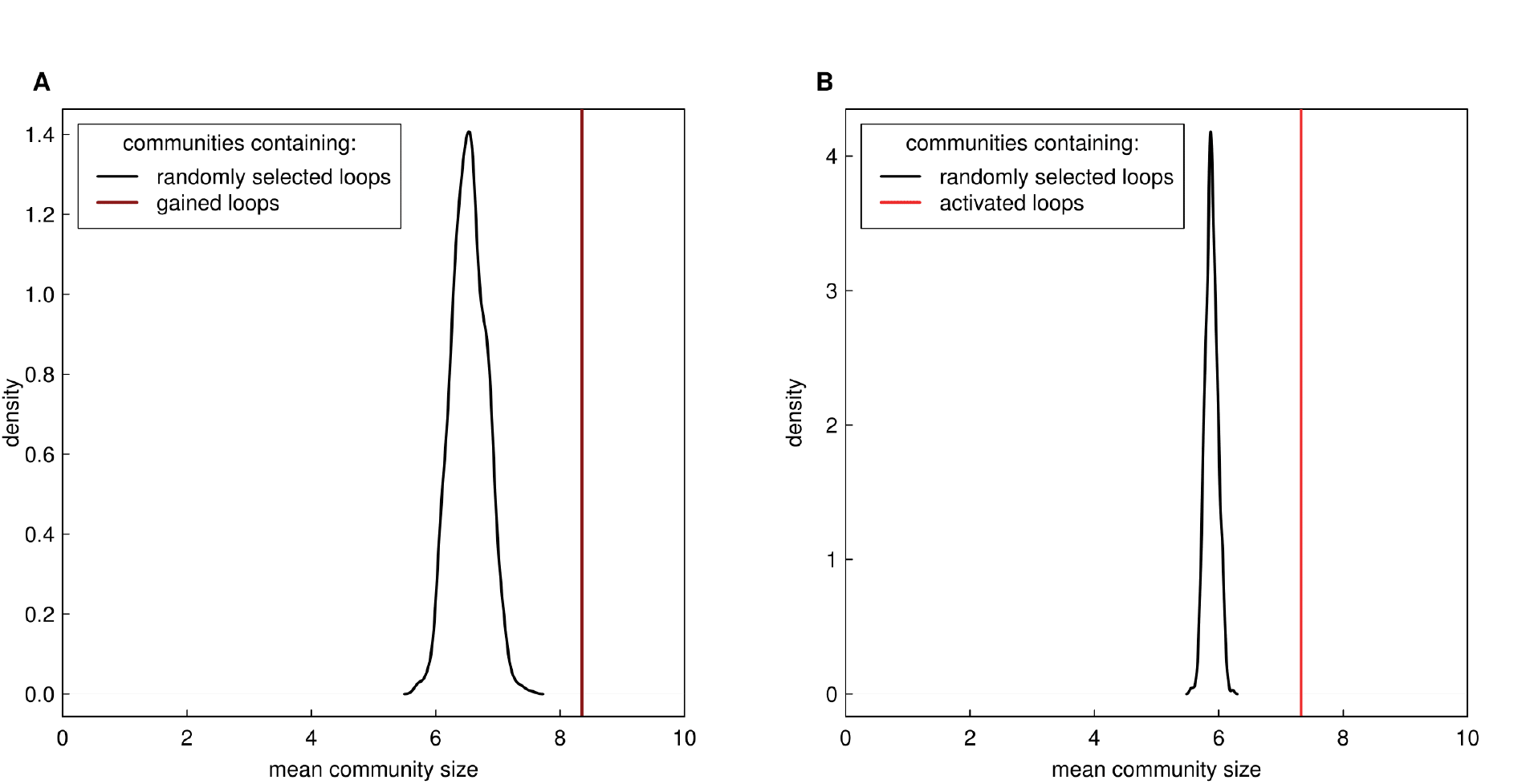
Community sizes of gained and activated loops compared to randomly chosen loops, Related to Figure 4. (**A**) 1000 random sets of loops, equal in number to gained loops, were selected. For each set we determined the mean size of communities that contained at least one selected loop. The distributions are shown in panel A. The mean community size for the observed gained loops is shown as a dark red line. (**B**) The same analysis was performed for activated loops.

**Figure S4:**
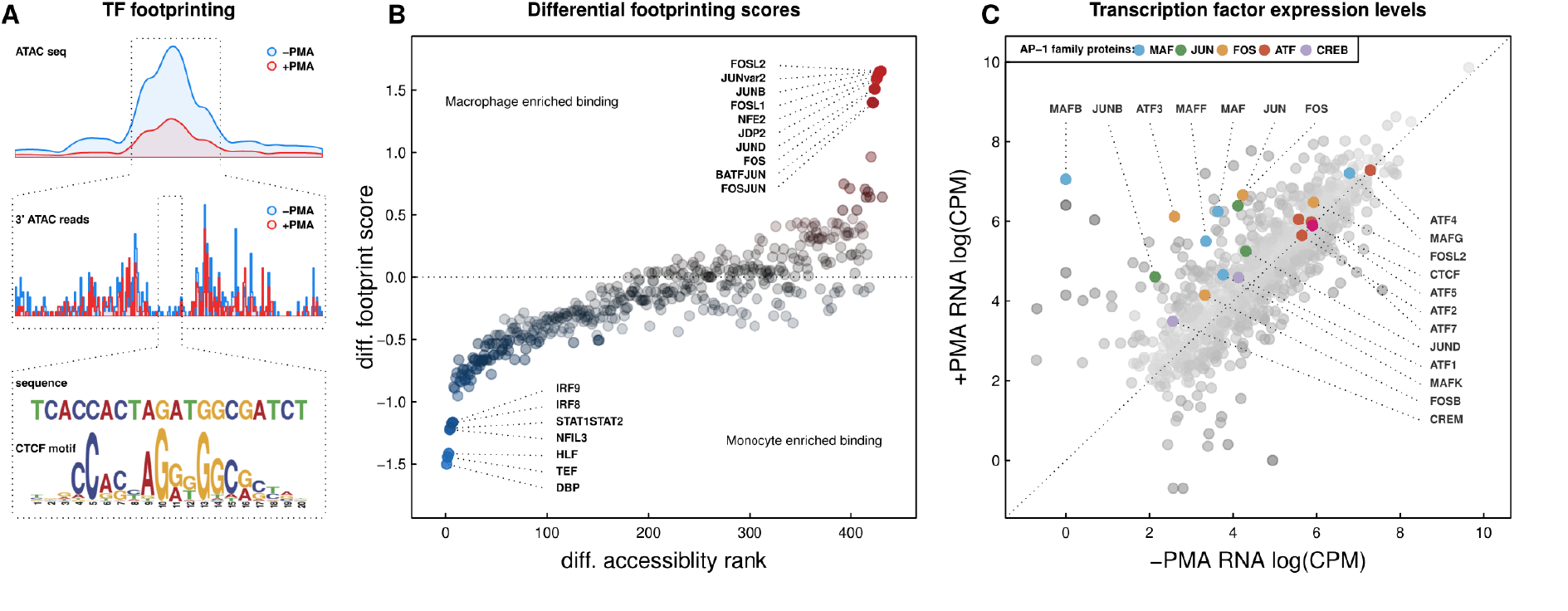
Increased expression and chromatin binding of AP-1 family genes during PMA-induced differentiation of THP-1 cells, Related to Figure 5. (**A**) Depiction of footprinting method. ATAC-seq signal track identifies accessible region (Top). Signal track depicting 3’ of ATAC seq reads reveals TF footprint (Middle). Sequence analysis within the footprint reveals CTCF motif (Bottom) (**B**) TF footprints plotted by differential median accessibility rank on the x-axis and median differential footprinting score on the y-axis. Both scores were determined by Wellington. (**C**) Scatter plot depicting gene level FPKM values in untreated and PMA-treated macrophages for all transcription factors. AP-1 family proteins and CTCF are colored.

**Figure S5:**
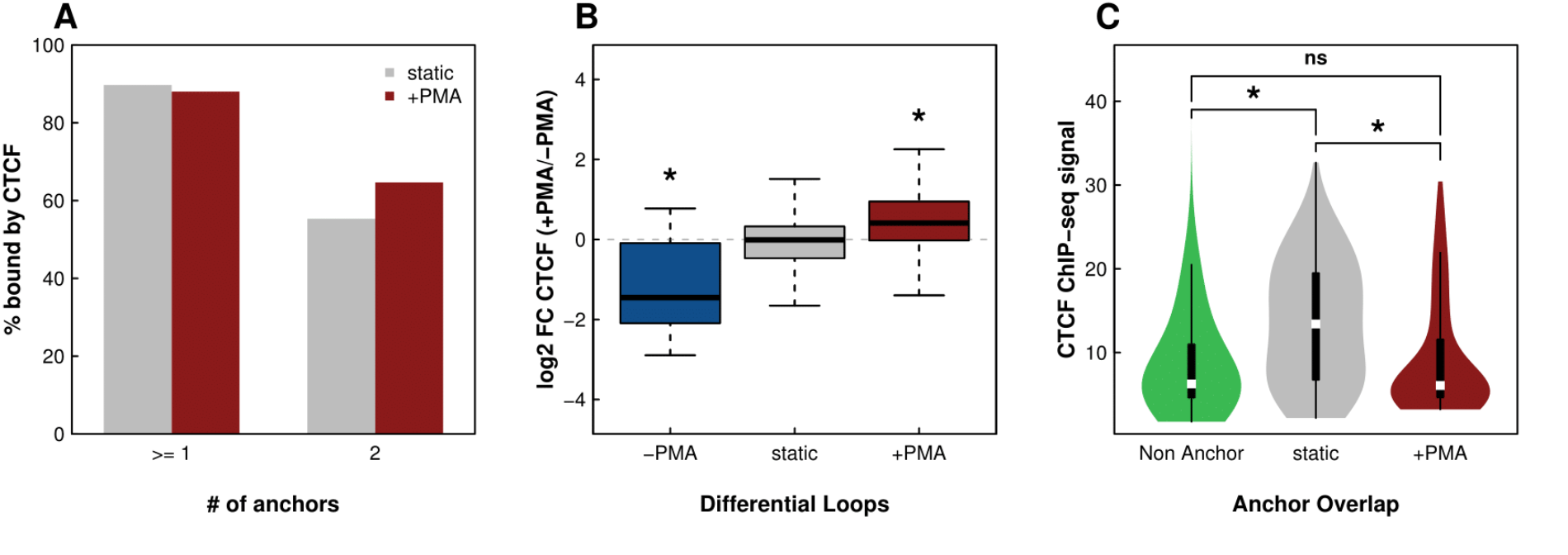
CTCF binds weakly at gained loop anchors, Related to Figure 5. (**A**) Bar plot depicting the percent of static (grey) or gained (red) loops with CTCF peaks at the anchors. No significant difference is detected between static and +PMA loops (p > 0.01 based on Fisher’s Exact Test). (**B**) Box plot showing the fold changes of CTCF acetylation peaks at lost, static, and gained loop anchors. Asterisks indicate p < 10^−3^ based on Wilcox Rank Sum Test. (**C**) Violin plot representing distributions of CTCF ChIP-seq signal at static, gained, and non-loop anchors. Asterisk indicate p < 10^−9^ based on Wilcox Rank Sum Test.

**Figure S6:**
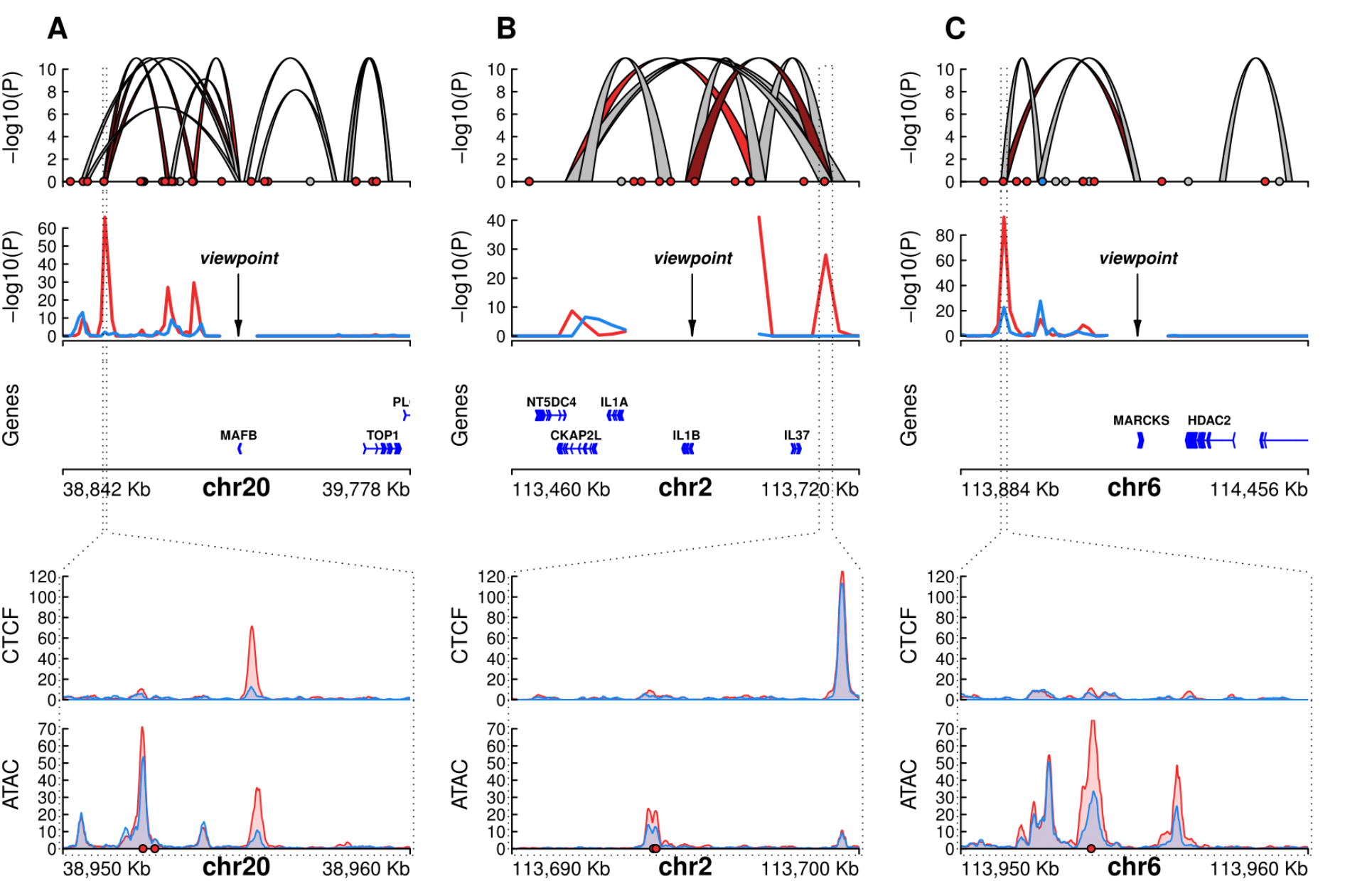
AP-1-bound activation hubs, Related to Figure 6. Three example regions harboring an AP-1-bound activation hub are shown (**A-C**). From top to bottom: DNA loops determined by in situ Hi-C are colored according to loop subset; −log10(P) of loop pixel enrichment compared to local background of loops connecting to the viewpoint indicated by an arrow (untreated shown in blue; PMA-treated shown in red); Genes and orientations depicted as arrows; CTCF binding profile at +PMA-specific anchor region; ATAC-seq signal at +PMA-specific anchor region. Top and bottom plots including circles indication presence of AP-1 footprint which are colored according to cell type specificity (red = increased in PMA-treated cells, blue = increased in non-treated cells, grey = no significant change).

## REFERENCES

Alexander, R.P., Fang, G., Rozowsky, J., Snyder, M., and Gerstein, M.B. (2010). Annotating noncoding regions of the genome. Nat Rev Genet 11, 559–571.

Beagan, J.A., Gilgenast, T.G., Kim, J., Plona, Z., Norton, H.K., Hu, G., Hsu, S.C., Shields, E.J., Lyu, X., Apostolou, E., et al. (2016). Local Genome Topology Can Exhibit an Incompletely Rewired 3D-Folding State during Somatic Cell Reprogramming. Cell Stem Cell 18, 611–624.

Boyle, A.P., Davis, S., Shulha, H.P., Meltzer, P., Margulies, E.H., Weng, Z., Furey, T.S., and Crawford, G.E. (2008). High-resolution mapping and characterization of open chromatin across the genome. Cell 132, 311–322.

Buenrostro, J.D., Wu, B., Chang, H.Y., and Greenleaf, W.J. (2015). ATAC-seq: A Method for Assaying Chromatin Accessibility Genome-Wide. Curr Protoc Mol Biol 109, 21 29 21–29.

Busslinger, G.A., Stocsits, R.R., van der Lelij, P., Axelsson, E., Tedeschi, A., Galjart, N., and Peters, J.M. (2017). Cohesin is positioned in mammalian genomes by transcription, CTCF and Wapl. Nature 544, 503–507.

Chavanas, S., Adoue, V., Mechin, M.C., Ying, S., Dong, S., Duplan, H., Charveron, M., Takahara, H., Serre, G., and Simon, M. (2008). Long-range enhancer associated with chromatin looping allows AP-1 regulation of the peptidylarginine deiminase 3 gene in differentiated keratinocyte. PLoS One 3, e3408.

Chawla, A., Nguyen, K.D., and Goh, Y.P. (2011). Macrophage-mediated inflammation in metabolic disease. Nat Rev Immunol 11, 738–749.

Clauset, A., Newman, M.E., and Moore, C. (2004). Finding community structure in very large networks. Phys Rev E Stat Nonlin Soft Matter Phys 70, 066111.

Creyghton, M.P., Cheng, A.W., Welstead, G.G., Kooistra, T., Carey, B.W., Steine, E.J., Hanna, J., Lodato, M.A., Frampton, G.M., Sharp, P.A., et al. (2010). Histone H3K27ac separates active from poised enhancers and predicts developmental state. Proc Natl Acad Sci U S A 107, 21931–21936.

Daigneault, M., Preston, J.A., Marriott, H.M., Whyte, M.K., and Dockrell, D.H. (2010). The identification of markers of macrophage differentiation in PMA-stimulated THP-1 cells and monocyte-derived macrophages. PLoS One 5, e8668.

Darrow, E.M., Huntley, M.H., Dudchenko, O., Stamenova, E.K., Durand, N.C., Sun, Z., Huang, S.C., Sanborn, A.L., Machol, I., Shamim, M., et al. (2016). Deletion of DXZ4 on the human inactive X chromosome alters higher-order genome architecture. Proc Natl Acad Sci U S A 113, E4504–4512.

Davies, J.O., Oudelaar, A.M., Higgs, D.R., and Hughes, J.R. (2017). How best to identify chromosomal interactions: a comparison of approaches. Nat Methods 14, 125–134.

Dekker, J., Marti-Renom, M.A., and Mirny, L.A. (2013). Exploring the three-dimensional organization of genomes: interpreting chromatin interaction data. Nat Rev Genet 14, 390–403.

Deng, W., Lee, J., Wang, H., Miller, J., Reik, A., Gregory, P.D., Dean, A., and Blobel, G.A. (2012). Controlling long-range genomic interactions at a native locus by targeted tethering of a looping factor. Cell 149, 1233–1244.

Deng, W., Rupon, J.W., Krivega, I., Breda, L., Motta, I., Jahn, K.S., Reik, A., Gregory, P.D., Rivella, S., Dean, A., et al. (2014). Reactivation of developmentally silenced globin genes by forced chromatin looping. Cell 158, 849–860.

Dixon, J.R., Jung, I., Selvaraj, S., Shen, Y., Antosiewicz-Bourget, J.E., Lee, A.Y., Ye, Z., Kim, A., Rajagopal, N., Xie, W., et al. (2015). Chromatin architecture reorganization during stem cell differentiation. Nature 518, 331–336.

Dixon, J.R., Selvaraj, S., Yue, F., Kim, A., Li, Y., Shen, Y., Hu, M., Liu, J.S., and Ren, B. (2012). Topological domains in mammalian genomes identified by analysis of chromatin interactions. Nature 485, 376–380.

Dostie, J., Richmond, T.A., Arnaout, R.A., Selzer, R.R., Lee, W.L., Honan, T.A., Rubio, E.D., Krumm, A., Lamb, J., Nusbaum, C., et al. (2006). Chromosome Conformation Capture Carbon Copy (5C): a massively parallel solution for mapping interactions between genomic elements. Genome Res 16, 1299–1309.

Dowen, J.M., Fan, Z.P., Hnisz, D., Ren, G., Abraham, B.J., Zhang, L.N., Weintraub, A.S., Schuijers, J., Lee, T.I., Zhao, K., et al. (2014). Control of cell identity genes occurs in insulated neighborhoods in mammalian chromosomes. Cell 159, 374–387.

Drissen, R., Palstra, R.J., Gillemans, N., Splinter, E., Grosveld, F., Philipsen, S., and de Laat, W. (2004). The active spatial organization of the beta-globin locus requires the transcription factor EKLF. Genes Dev 18, 2485–2490.

Durand, N.C., Shamim, M.S., Machol, I., Rao, S.S., Huntley, M.H., Lander, E.S., and Aiden, E.L. (2016). Juicer Provides a One-Click System for Analyzing Loop-Resolution Hi-C Experiments. Cell Syst 3, 95–98.

Eferl, R., and Wagner, E.F. (2003). AP-1: a double-edged sword in tumorigenesis. Nat Rev Cancer 3, 859–868.

Fullwood, M.J., Liu, M.H., Pan, Y.F., Liu, J., Xu, H., Mohamed, Y.B., Orlov, Y.L., Velkov, S., Ho, A., Mei, P.H., et al. (2009). An oestrogen-receptor-alpha-bound human chromatin interactome. Nature 462, 58–64.

Giresi, P.G., Kim, J., McDaniell, R.M., Iyer, V.R., and Lieb, J.D. (2007). FAIRE (Formaldehyde-Assisted Isolation of Regulatory Elements) isolates active regulatory elements from human chromatin. Genome Res 17, 877–885.

Guo, Y., Xu, Q., Canzio, D., Shou, J., Li, J., Gorkin, D.U., Jung, I., Wu, H., Zhai, Y., Tang, Y., et al. (2015). CRISPR Inversion of CTCF Sites Alters Genome Topology and Enhancer/Promoter Function. Cell 162, 900–910.

Haarhuis, J.H.I., van der Weide, R.H., Blomen, V.A., Yanez-Cuna, J.O., Amendola, M., van Ruiten, M.S., Krijger, P.H.L., Teunissen, H., Medema, R.H., van Steensel, B., et al. (2017). The Cohesin Release Factor WAPL Restricts Chromatin Loop Extension. Cell 169, 693–707 e614.

Heidari, N., Phanstiel, D.H., He, C., Grubert, F., Jahanbani, F., Kasowski, M., Zhang, M.Q., and Snyder, M.P. (2014). Genome-wide map of regulatory interactions in the human genome. Genome Res 24, 1905–1917.

Hnisz, D., Weintraub, A.S., Day, D.S., Valton, A.L., Bak, R.O., Li, C.H., Goldmann, J., Lajoie, B. R., Fan, Z.P., Sigova, A.A., et al. (2016). Activation of proto-oncogenes by disruption of chromosome neighborhoods. Science 351, 1454–1458.

Imakaev, M., Fudenberg, G., McCord, R.P., Naumova, N., Goloborodko, A., Lajoie, B.R., Dekker, J., and Mirny, L.A. (2012). Iterative correction of Hi-C data reveals hallmarks of chromosome organization. Nat Methods 9, 999–1003.

Jin, F., Li, Y., Dixon, J.R., Selvaraj, S., Ye, Z., Lee, A.Y., Yen, C.A., Schmitt, A.D., Espinoza, C.A., and Ren, B. (2013). A high-resolution map of the three-dimensional chromatin interactome in human cells. Nature 503, 290–294.

Kelly, L.M., Englmeier, U., Lafon, I., Sieweke, M.H., and Graf, T. (2000). MafB is an inducer of monocytic differentiation. EMBO J 19, 1987–1997.

Kieffer-Kwon, K.R., Tang, Z., Mathe, E., Qian, J., Sung, M.H., Li, G., Resch, W., Baek, S., Pruett, N., Grontved, L., et al. (2013). Interactome maps of mouse gene regulatory domains reveal basic principles of transcriptional regulation. Cell 155, 1507–1520.

Kim, A., and Dean, A. (2012). Chromatin loop formation in the beta-globin locus and its role in globin gene transcription. Mol Cells 34, 1–5.

Knight, P.A., and Ruiz, D. (2013). A fast algorithm for matrix balancing. Ima Journal of Numerical Analysis 33, 1029–1047.

Kouno, T., de Hoon, M., Mar, J.C., Tomaru, Y., Kawano, M., Carninci, P., Suzuki, H., Hayashizaki, Y., and Shin, J.W. (2013). Temporal dynamics and transcriptional control using single-cell gene expression analysis. Genome Biol 14, R118.

Krijger, P.H., and de Laat, W. (2016). Regulation of disease-associated gene expression in the 3D genome. Nat Rev Mol Cell Biol 17, 771–782.

Krivega, I., and Dean, A. (2016). Chromatin looping as a target for altering erythroid gene expression. Ann N Y Acad Sci 1368, 31–39.

Lieberman-Aiden, E., van Berkum, N.L., Williams, L., Imakaev, M., Ragoczy, T., Telling, A., Amit, I., Lajoie, B.R., Sabo, P.J., Dorschner, M.O., et al. (2009). Comprehensive mapping of long-range interactions reveals folding principles of the human genome. Science 326, 289–293.

Liebermann, D.A., Gregory, B., and Hoffman, B. (1998). AP-1 (Fos/Jun) transcription factors in hematopoietic differentiation and apoptosis. Int J Oncol 12, 685–700.

Mechta-Grigoriou, F., Gerald, D., and Yaniv, M. (2001). The mammalian Jun proteins: redundancy and specificity. Oncogene 20, 2378–2389.

Merkenschlager, M., and Nora, E.P. (2016). CTCF and Cohesin in Genome Folding and Transcriptional Gene Regulation. Annu Rev Genomics Hum Genet 17, 17–43.

Montavon, T., Soshnikova, N., Mascrez, B., Joye, E., Thevenet, L., Splinter, E., de Laat, W., Spitz, F., and Duboule, D. (2011). A regulatory archipelago controls Hox genes transcription in digits. Cell 147, 1132–1145.

Moore, K.J., Sheedy, F.J., and Fisher, E.A. (2013). Macrophages in atherosclerosis: a dynamic balance. Nat Rev Immunol 13, 709–721.

Mumbach, M.R., Rubin, A.J., Flynn, R.A., Dai, C., Khavari, P.A., Greenleaf, W.J., and Chang, H.Y. (2016). HiChIP: efficient and sensitive analysis of protein-directed genome architecture. Nat Methods 13, 919–922.

Narendra, V., Bulajic, M., Dekker, J., Mazzoni, E.O., and Reinberg, D. (2016). CTCF-mediated topological boundaries during development foster appropriate gene regulation. Genes Dev 30, 2657–2662.

Nora, E.P., Lajoie, B.R., Schulz, E.G., Giorgetti, L., Okamoto, I., Servant, N., Piolot, T., van Berkum, N.L., Meisig, J., Sedat, J., et al. (2012). Spatial partitioning of the regulatory landscape of the X-inactivation centre. Nature 485, 381–385.

Noy, R., and Pollard, J.W. (2014). Tumor-associated macrophages: from mechanisms to therapy. Immunity 41, 49–61.

Odero, M.D., Zeleznik-Le, N.J., Chinwalla, V., and Rowley, J.D. (2000). Cytogenetic and molecular analysis of the acute monocytic leukemia cell line THP-1 with an MLL-AF9 translocation. Genes Chromosomes Cancer 29, 333–338.

Ong, C.T., and Corces, V.G. (2014). CTCF: an architectural protein bridging genome topology and function. Nat Rev Genet 15, 234–246.

Ostuni, R., Kratochvill, F., Murray, P.J., and Natoli, G. (2015). Macrophages and cancer: from mechanisms to therapeutic implications. Trends Immunol 36, 229–239.

Palstra, R.J., Tolhuis, B., Splinter, E., Nijmeijer, R., Grosveld, F., and de Laat, W. (2003). The beta-globin nuclear compartment in development and erythroid differentiation. Nat Genet 35, 190–194.

Phillips-Cremins, J.E., Sauria, M.E., Sanyal, A., Gerasimova, T.I., Lajoie, B.R., Bell, J.S., Ong, C.T., Hookway, T.A., Guo, C., Sun, Y., et al. (2013). Architectural protein subclasses shape 3D organization of genomes during lineage commitment. Cell 153, 1281–1295.

Piper, J., Assi, S.A., Cauchy, P., Ladroue, C., Cockerill, P.N., Bonifer, C., and Ott, S. (2015). Wellington-bootstrap: differential DNase-seq footprinting identifies cell-type determining transcription factors. BMC Genomics 16, 1000.

Qiao, Y., Shiue, C.N., Zhu, J., Zhuang, T., Jonsson, P., Wright, A.P., Zhao, C., and Dahlman-Wright, K. (2015). AP-1-mediated chromatin looping regulates ZEB2 transcription: new insights into TNFalpha-induced epithelial-mesenchymal transition in triple-negative breast cancer. Oncotarget 6, 7804–7814.

Quinlan, A.R., and Hall, I.M. (2010). BEDTools: a flexible suite of utilities for comparing genomic features. Bioinformatics 26, 841–842.

Rao, S.S., Huntley, M.H., Durand, N.C., Stamenova, E.K., Bochkov, I.D., Robinson, J.T.,

Sanborn, A.L., Machol, I., Omer, A.D., Lander, E.S., et al. (2014). A 3D map of the human genome at kilobase resolution reveals principles of chromatin looping. Cell 159, 1665–1680.

Ritchie, M.E., Phipson, B., Wu, D., Hu, Y., Law, C.W., Shi, W., and Smyth, G.K. (2015). limma powers differential expression analyses for RNA-sequencing and microarray studies. Nucleic Acids Res 43, e47.

Roadmap Epigenomics, C., Kundaje, A., Meuleman, W., Ernst, J., Bilenky, M., Yen, A., Heravi-Moussavi, A., Kheradpour, P., Zhang, Z., Wang, J., et al. (2015). Integrative analysis of 111 reference human epigenomes. Nature 518, 317–330.

Rosa, A., Ballarino, M., Sorrentino, A., Sthandier, O., De Angelis, F.G., Marchioni, M., Masella, B., Guarini, A., Fatica, A., Peschle, C., et al. (2007). The interplay between the master transcription factor PU.1 and miR-424 regulates human monocyte/macrophage differentiation. Proc Natl Acad Sci U S A 104, 19849–19854.

Sanyal, A., Lajoie, B.R., Jain, G., and Dekker, J. (2012). The long-range interaction landscape of gene promoters. Nature 489, 109–113.

Schmitt, A.D., Hu, M., Jung, I., Xu, Z., Qiu, Y., Tan, C.L., Li, Y., Lin, S., Lin, Y., Barr, C.L., et al. (2016). A Compendium of Chromatin Contact Maps Reveals Spatially Active Regions in the Human Genome. Cell Rep 17, 2042–2059.

Shaulian, E., and Karin, M. (2002). AP-1 as a regulator of cell life and death. Nat Cell Biol 4, E131–136.

Sherwood, R.I., Hashimoto, T., O'Donnell, C.W., Lewis, S., Barkal, A.A., van Hoff, J.P., Karun, V., Jaakkola, T., and Gifford, D.K. (2014). Discovery of directional and nondirectional pioneer transcription factors by modeling DNase profile magnitude and shape. Nat Biotechnol 32, 171–178.

Smith, E.M., Lajoie, B.R., Jain, G., and Dekker, J. (2016). Invariant TAD Boundaries Constrain Cell-Type-Specific Looping Interactions between Promoters and Distal Elements around the CFTR Locus. Am J Hum Genet 98, 185–201.

Sofueva, S., Yaffe, E., Chan, W.C., Georgopoulou, D., Vietri Rudan, M., Mira-Bontenbal, H., Pollard, S.M., Schroth, G.P., Tanay, A., and Hadjur, S. (2013). Cohesin-mediated interactions organize chromosomal domain architecture. EMBO J 32, 3119–3129.

Song, S.H., Hou, C., and Dean, A. (2007). A positive role for NLI/Ldb1 in long-range beta-globin locus control region function. Mol Cell 28, 810–822.

Tolhuis, B., Palstra, R.J., Splinter, E., Grosveld, F., and de Laat, W. (2002). Looping and interaction between hypersensitive sites in the active beta-globin locus. Mol Cell 10, 1453–1465.

Vakoc, C.R., Letting, D.L., Gheldof, N., Sawado, T., Bender, M.A., Groudine, M., Weiss, M.J., Dekker, J., and Blobel, G.A. (2005). Proximity among distant regulatory elements at the betaglobin locus requires GATA-1 and FOG-1. Mol Cell 17, 453–462.

Valledor, A.F., Borras, F.E., Cullell-Young, M., and Celada, A. (1998). Transcription factors that regulate monocyte/macrophage differentiation. J Leukoc Biol 63, 405–417.

Wang, J., Zhuang, J., Iyer, S., Lin, X.Y., Greven, M.C., Kim, B.H., Moore, J., Pierce, B.G., Dong, X., Virgil, D., et al. (2013). Factorbook.org: a Wiki-based database for transcription factor-binding data generated by the ENCODE consortium. Nucleic Acids Res 41, D171–176.

